# Dynamic model of carbon dioxide-induced stomatal closure reveals a feedback core for cellular decision-making

**DOI:** 10.1101/2023.10.23.563079

**Authors:** Xiao Gan, Palanivelu Sengottaiyan, Kyu Hyong Park, Sarah M. Assmann, Réka Albert

## Abstract

Stomata are pores on plant aerial surfaces, each bordered by a pair of guard cells. They control gas exchange vital for plant survival. Understanding how guard cells respond to environmental signals such as atmospheric carbon dioxide (CO_2_) levels is not only insightful to fundamental biology but also relevant to real-world issues of crop productivity under global climate change. In the past decade, multiple important signaling elements for stomatal closure induced by elevated CO_2_ have been identified. Yet, there is no comprehensive understanding of high CO_2_ induced stomatal closure. In this work we assemble a cellular signaling network underlying high CO_2_-induced stomatal closure by integrating evidence from a comprehensive literature analysis. We further construct a Boolean dynamic model of the network, which allows *in silico* simulation of the stomatal closure response to high CO_2_ in wild-type *Arabidopsis thaliana* plants and in cases of pharmacological or genetic manipulation of network nodes. Our model has a 91% accuracy in capturing known experimental observations. We perform network-based logical analysis and reveal a feedback core of the network, which dictates cellular decisions in closure response to high CO_2_. Based on these analyses, we predict and experimentally confirm that applying nitric oxide (NO) induces stomatal closure in ambient CO_2_ and causes hypersensitivity to elevated CO_2_. Moreover, we predict a negative regulatory relationship between NO and the protein phosphatase ABI2 and find experimentally that NO inhibits ABI2 phosphatase activity. The experimental validation of these model predictions demonstrates the effectiveness of network-based modeling and highlights the decision-making role of the feedback core of the network in signal transduction. We further explore the model’s potential in predicting targets of signaling elements not yet connected to the CO_2_ network. Our combination of network science, *in silico* model simulation, and experimental assays demonstrates an effective interdisciplinary approach to understanding system-level biology.

## I. Introduction

Stomata are microscopic pores on plant leaf surfaces that control gas exchange such as the uptake of CO_2_, and the release of water vapor and photosynthetically-produced O_2_. Each stomate is bordered by two guard cells, whose shape change in response to the external and internal environment controls the stomate’s opening or closure. Because plants inevitably lose water via transpiration through the stomata, the size of the stomatal pores is tightly regulated to balance the competing needs of CO_2_ supply for photosynthesis and plant hydration. Stomata open in response to environmental signals such as light and low atmospheric CO_2_ levels and close in response to darkness, high CO_2_, drought, and low humidity [1]. As elucidated in detail for the stress hormone abscisic acid (ABA) [2–4], stimulus perception activates an intracellular signal transduction network, which leads to Ca^2+^ influx and anion efflux, with consequent membrane depolarization promoting K^+^ efflux. The outflow of ions increases cellular water potential, creating a driving force for water efflux through aquaporins, which finally causes guard cell deflation and stomatal closure. However, the extent to which the ABA signal transduction network is shared by other closure signals such as darkness and high CO_2_ is currently not elucidated [2].

Understanding how stomata respond to high CO_2_ is a fundamental question for plant biology. Since stomata control gas exchange vital for plant water retention and photosynthesis, stomatal response to abiotic stress is an essential process for plant survival. Furthermore, understanding plant response to high CO_2_ also has imperative practical significance: over the past 60 years, CO_2_ concentration in the atmosphere has increased by 32% (NOAA at Mauna Loa [5]). The greenhouse effect induced by increased CO_2_ has increased global temperature, which exacerbates drought, compounding the stress. Plant stomatal responses to CO_2_ are therefore highly relevant to real-world issues of crop productivity under the adversity posed by climate change.

### Current knowledge of high CO_2_ signaling in guard cells

Stomatal movement in response to high CO_2_ concentration is a complex process in which numerous cellular elements have been implicated by prior studies. These elements interact with and regulate each other via chemical reactions, protein interactions, and physical processes, forming a complex cell signaling network that propagates the signal from CO_2_ to stomatal closure. Atmospheric CO_2_ enters guard cells through CO_2_ porins [6]. The CO_2_-binding carbonic anhydrase enzymes CA1 and CA4 catalyze the reversible chemical reaction CO_2_ + H_2_O → HCO_3_^−^ + H^+^, and these enzymes are a demonstrated component of the CO_2_ signal transduction mechanism [7, 8]. HT1 (high leaf temperature 1) is an important CO_2_ signaling element that inhibits the activation of the main anion channel SLAC1 (SLOW ANION CHANNEL-ASSOCIATED 1). Multiple mutants of HT1 have been shown to result in insensitivity to high CO_2_ [9–11]. Mitogen-activated protein kinases 4 and 12 (MPK4/12) bind to HT1 and inhibit its kinase activity [11, 12]. Bicarbonate enhances and stabilizes the binding of MPK4/12 to HT1 [13, 14]. Arabidopsis RESISTANT TO HIGH CO_2_ (RHC1), a MATE-type transporter, may also link elevated CO_2_ concentration to repression of HT1, although this has been debated [15, 16]. Disruption of the second messengers Ca^2+^, ROS (reactive oxygen species), or NO (nitric oxide) impairs high CO_2_ induced closure [17–22], implicating them in CO_2_ signaling. Despite the recent identification of key elements and pathways in the high CO_2_ response, further elucidation is needed of how the known elements and pathways interact with each other to induce stomatal closure. A systems biology approach integrating existing evidence into a signaling network can identify the gaps of knowledge and suggest potential ways to elucidate all the mechanisms that contribute to stomatal closure.

### Prior modeling of guard cell signaling networks

In the past, our team has developed multiple networks and network-based models to characterize the guard cell signaling system. In 2006, Li et al. constructed a Boolean model of ABA (abscisic acid)-induced stomatal closure [23]. In 2014, Sun et al. developed a multi-level discrete dynamic model focusing on light-induced stomatal opening [24]. In 2017, Albert et al. made major updates to the ABA-induced stomatal closure model, including many more signaling elements discovered in recent years [3]. These network-based models revealed mechanistic insights and made novel predictions that were validated by follow-up experiments. Furthermore, we have developed novel theoretical concepts such as the concept of the stable motif, which forms the foundation of computational methods to determine the repertoire of long-time behaviors of biological systems and to identify nodes that can drive the system into a desired state [25–28].

Karanam et al. recently developed a graphical user interface to easily analyze Boolean models [29] and applied it to integrate a CO_2_ signaling pathway into our previous ABA-induced closure model [3], as an approach to describe CO_2_ signaling. While it is a good demonstration of their methodology, this preliminary model is not a rigorous representation of CO_2_ signaling in that: (1) it lacks a comprehensive literature review to identify elements that are differentially regulated in high CO_2_ signaling as compared to ABA signaling; (2) it assumes that high CO_2_ is sufficient to directly activate water efflux through the membrane, as well as Ca^2+^ inflow, which together are sufficient to drive stomatal closure: these assumptions short-circuit all other CO_2_-related nodes, i.e. render the rest of their network superfluous. Because of the assumed direct connection to water efflux, this model would fail to recapitulate, for example, the experimental observations of impaired high CO_2_ induced closure for *ht1* [10] and *mpk4/12* [12] mutants. A more rigorous network construction approach is required to produce a more accurate representation and simulation of high CO_2_ induced closure.

In this work, we compile and integrate an extensive body of literature evidence to assemble a guard cell signal transduction network that characterizes how high CO_2_ induces stomatal closure. To further capture the dynamics involved in the stomatal closure process, we develop a dynamic model of the signaling network. We rigorously evaluate the accuracy of the model against known experimental observations, obtaining a high (91%) accuracy. We perform analyses including network connectivity, model simulation and stable motif analysis, which reveal a feedback core that dictates the cellular decisions in the CO_2_ response, and the roles of key nodes in the process. We then demonstrate the predictive power of our network model by predicting the outcomes of numerous perturbations (e.g., node knockouts) that can potentially influence stomatal response; such predictions are valuable for the prioritization of wet bench experiments. Finally, we experimentally test and validate three predictions related to the role of NO in stomatal closure. A flowchart of our work is shown below (Figure 1). Our modeling framework captures system-level biological processes and provides insights into their logic and key underlying mechanisms.

**Figure 1.**
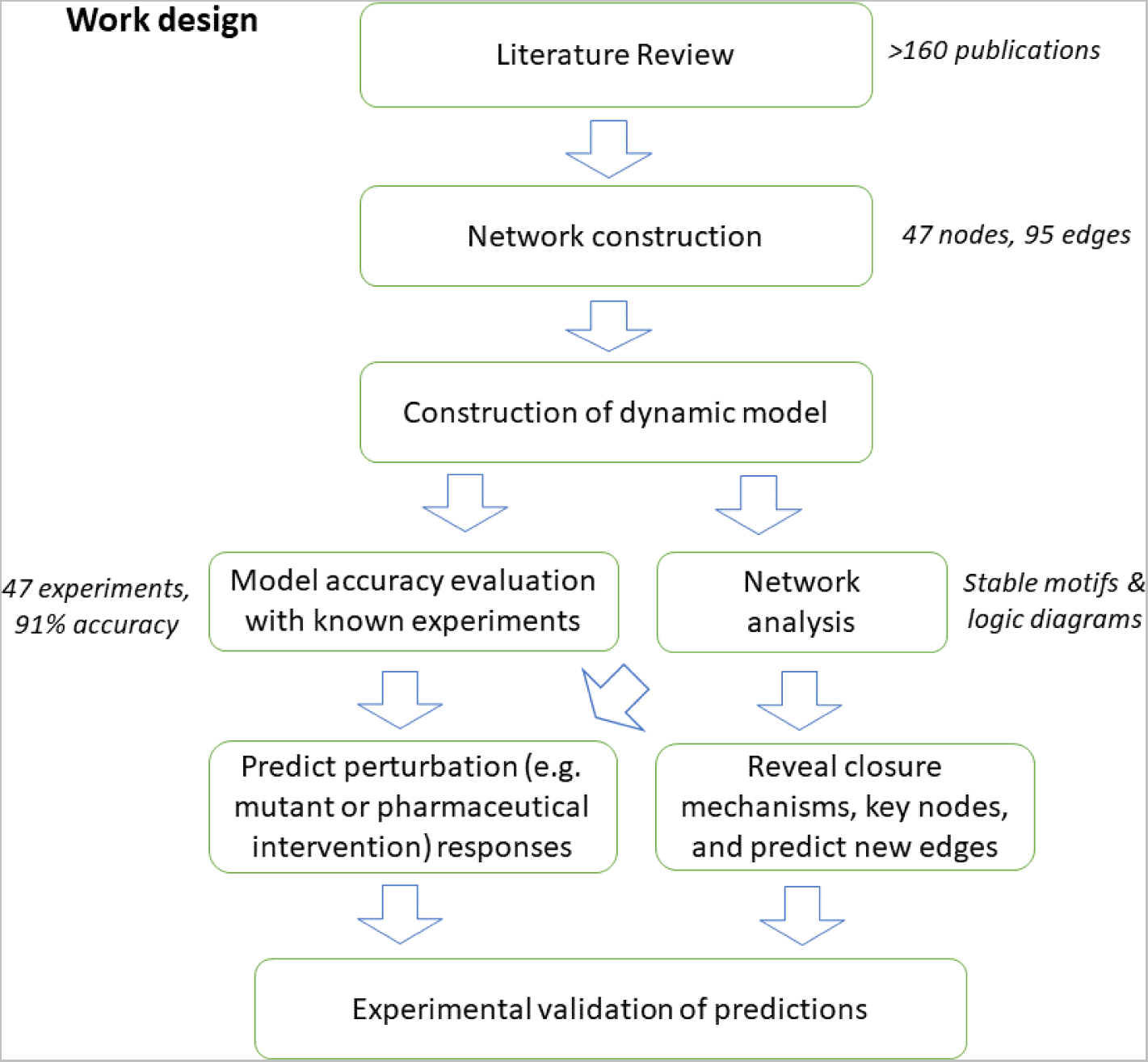
Flow chart of the overall work design. We construct a signaling network based on literature review, resulting in a network of 47 nodes and 95 edges. We then construct a Boolean dynamic model, which adds node states and regulatory logic to the network. We assess the accuracy of our network model by comparing its results with known experimental observations, finding that the model simulations successfully recapitulate 91% of experimental observations. We also perform multiple network-based analyses on the model. Finally, we experimentally validate selected model predictions.

## II. Results

### Construction and analysis of the signaling network of high CO_2_-induced stomatal closure

We conducted a comprehensive literature review, covering more than 160 publications relating to stomatal closure or to elements of guard cell CO_2_ signaling, and manually curated the interaction information from these publications (Supporting Info S1). Manual curation was necessary and important because not all guard cell signaling elements are equally relevant to high CO_2_ induced closure. For example, cytosolic pH increases in response to ABA, but does not increase under high CO_2_ [8, 30, 31]. Another example is the kinase OST1 (*Open Stomata1*/SnRK2.6), known to phosphorylate SLAC1 as a key contributor to the ion flow that leads to stomatal closure induced by ABA. Although *ost1* mutants show strongly impaired stomatal closure under both ABA and high CO_2_, a recent study showed that unlike ABA, high CO_2_ does not increase OST1 kinase activity in guard cells [32, 33].

To focus on CO_2_ signaling, we divide the evidence into five categories based on relevance: (1) elements and their interactions that are validated to be involved in CO_2_ signaling; (2) generic interactions that can be assumed to be involved in CO_2_ signaling, e.g. chemical reactions whose reactants are reasonably expected to be present in the guard cell; (3) evidence that shows an element is not involved in CO_2_ signaling; (4) evidence that supports an element’s role in stomatal closure in response to another signal, but the element’s involvement in high CO_2_-induced closure has not been investigated; (5) evidence that supports an element’s role in high CO_2_ induced closure but this element has no or insufficient connections to elements in categories (1) and (2). The resulting list of observations and their categorization is provided in Supporting Info S1. We use category (1) and (2) evidence to construct the high CO_2_ signaling network and exclude category (3) and (4) evidence. Category (5) evidence is also necessarily excluded, as it is not possible to connect such nodes to the network. Such categorization is also helpful to wet bench investigators; e.g., category 4 and 5 elements would be of particular interest to investigate further in the future.

To represent signaling elements with different functions, we deploy a multi-node representation for the OST1, MPK and SLAC1 proteins, separating the different (e.g., CO_2_-dependent and independent) mechanisms for their activation (see Figure 2C for an example, and see Supporting Info S2 and S3 for details). As the follow-up analyses require that each edge of the network points to a node, during network construction we redirected each evidence of influence on an edge to an equivalent influence on the target of the edge [34].

**Figure 2.**
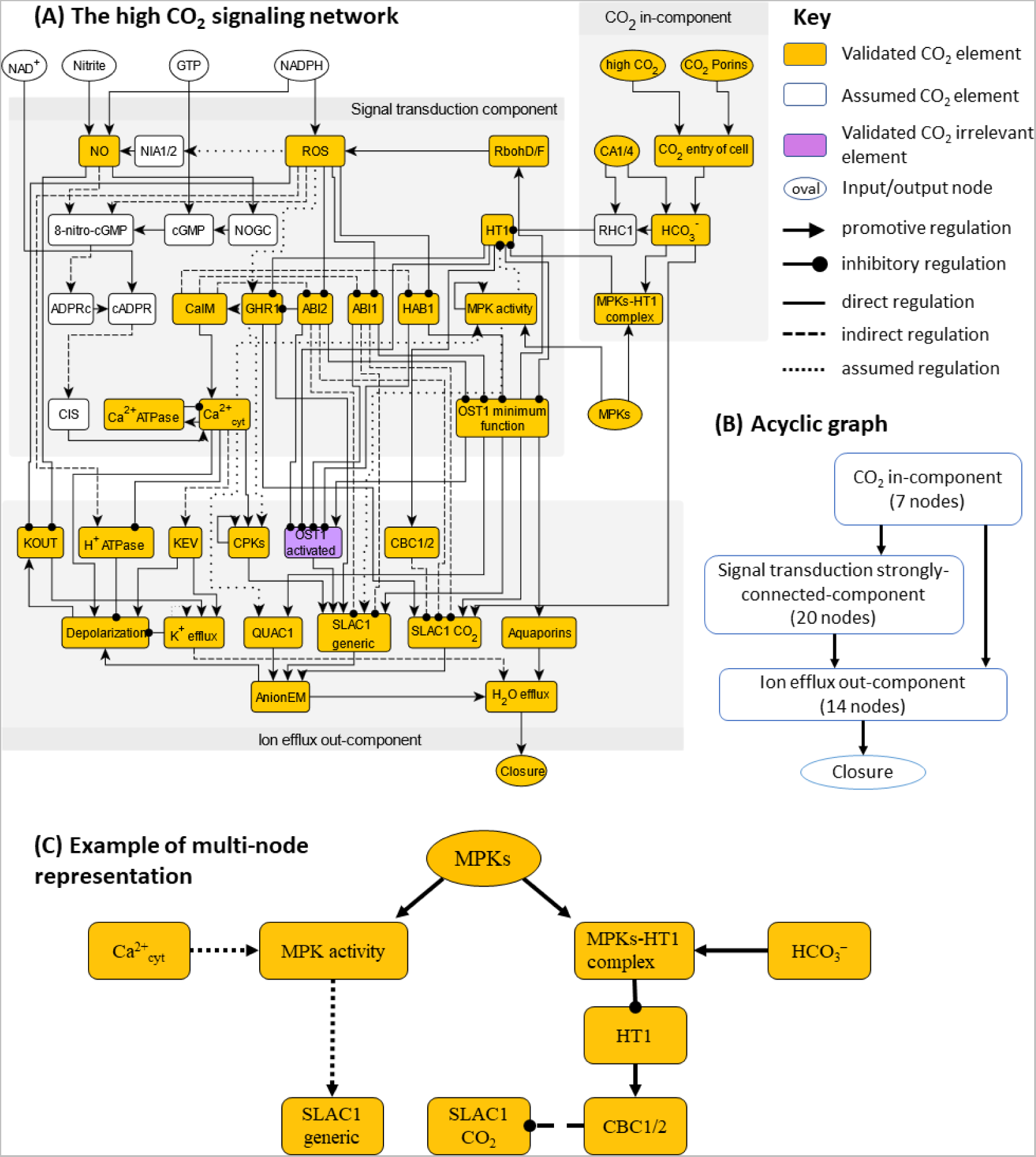
(A) The high CO_2_ signaling network with 47 nodes and 95 edges. (B) Acyclic graph representation of the network highlighting the in-component, the strongly-connected-component and the out-component, forming a hierarchical structure of signal transduction; (C) Illustration of the multi-node representation of signaling elements that participate in multiple pathways and mechanisms, using MPKs as an example. MPKs are known to inhibit HT1 by a physical interaction induced by bicarbonate and independent of the MPKs’ kinase activity [13, 14], and also are necessary in Ca^2+^-induced SLAC1 activation [35].

### Biological description of the high CO_2_-induced stomatal closure network and its overlap with the ABA-induced closure network

The signaling network corresponding to high CO_2_ induced closure contains 47 nodes and 95 edges (Figure 2A). A summary table of all edges is presented in Table 1. The full list of nodes and edges, including the full node names, is provided in Supporting Info S2. We mark nodes of the network (Figure 2A) in three colors, to reflect their specificity to high-CO_2_ induced closure. Yellow nodes represent validated elements in CO_2_ signaling, i.e., category (1); white nodes represent assumed elements based on generic evidence, i.e., category (2). Although by default nodes in categories (3) - (5) are not included in the network, “OST1 activated” is a member of a node pair (it represents increased kinase activity of OST1), thus we include it and mark it in purple to indicate that it belongs to category (4): it is not involved in high CO_2_-induced closure but is important under ABA-induced closure.

**Table 1.** Signaling elements that differ in the networks that lead to closure in response to ABA versus high CO_2_.

| Category | Nodes/signaling elements |
| --- | --- |
| CO<sub>2</sub>-specific signaling elements that are not involved in ABA-induced closure | CA1/4, HCO<sub>3</sub><sup>-</sup>, RHC1, HT1, MPKs-HT1 inhibition |
| ABA signaling elements observed to be absent or not significant in high CO<sub>2</sub>-induced closure | Cytosolic pH increase, OST1 activation |
| ABA signaling elements* whose participation in high CO<sub>2</sub> signaling has not been tested | AtRAC1, actin reorganization, GPA1, Malate, Microtubule depolymerization, PEPC, PP2CA, ROP11, S1P/PhytoS1P, SPHK1/2, SLAH3, TCTP;<br>*Phospholipases and molecules involved in reactions catalyzed by them:* PtdIns(4,5)P2, PLC, DAG, InsP3, InsP6, PLDα, PLDδ, PA;<br>*Elements of the pathway leading to vacuolar acidification:* PI3P5K, PtdIns(3,5)P2, V-PPase, V-ATPase, vacuolar acidification; |
| CO<sub>2</sub> signaling elements with insufficient regulatory information to integrate into our CO<sub>2</sub> network** | TPK1, KIN7, BIG1, AtABCB14, GCA2 |

\*The full names of the ABA signaling elements can be found in [3].
\*\* Details and references for these elements are provided in Supporting Info S1.

The edges in the network are also marked to distinguish their properties: solid edges indicate a direct interaction/regulation, dashed edges indicate indirect regulation, and dotted lines indicate assumed regulation. An example of indirect regulation is the edge from ROS to the H^+^ ATPase, which is dashed. An example of an edge referring to assumed regulation is the edge from HCO_3_^−^ to SLAC1 CO_2_, which is supported by a specific SLAC1 residue that is required for full CO_2_ signaling and is structurally predicted to interact with HCO_3_^−^ [36], but this predicted physical interaction has not been experimentally verified. Like this example, all assumed edges in the model are well-supported. Further justification of the assumed edges can be found in Supporting Info S3.

The high CO_2_-induced stomatal closure network is known to share nodes and pathways with the ABA-induced closure network, but it also has its unique signaling elements. Furthermore, through the literature review process, we also found multiple elements that participate in ABA-induced closure but are not yet evaluated in the context of high CO_2_ induced closure (also see Supporting Info S1 and S4). We list in Table 1 the different categories for the combined set of the signaling elements (i.e. nodes) of this CO_2_ network and of our 2017 ABA network [3].

### The CO_2_ signaling network consists of three distinct modules

Of the 47 nodes in the network, 8 (including high CO_2_) are source nodes that are not regulated by any other node, and one node (Closure) is a sink node (output). The source nodes other than the signal “high CO_2_” mainly represent molecules needed as substrates for reactions incorporated in the network: CO_2_ porins, CA1/4, GTP, MPKs, NAD^+^, NADPH, Nitrite. Structural and connectivity analysis shows that the CO_2_ network is organized into three graph modules (called “connected components” in graph theoretical language), as shown in Figure 2B. First, 7 early CO_2_ signaling nodes form an in-component, i.e., these nodes are not regulated by any nodes from other components in the network. This module mainly represents how CO_2_ enters the guard cell and is converted to HCO_3_^−^. Second, a large strongly-connected-component (SCC) of 20 nodes forms the core of the signal transduction network. “Strongly connected” means that for any node pair A, B, there is at least one regulatory path from A to B and at least one path from B to A. The feedback loops underlying the strong connectivity allows complex dynamical behavior that ultimately dictates the homeostasis of the system. The four edges between the in-component and the SCC represent CO_2_ sensing mechanisms, e.g. the inhibitory effect of HCO_3_^−^ on HT1 via the formation of the MPKs-HT1 complex. Third, 13 nodes, mainly representing ion transport mechanisms and the ion and water fluxes they mediate, as well as the output node “Closure”, form the out-component that ultimately implements stomatal closure. These nodes are mainly regulated by nodes of the SCC and do not regulate nodes in other connected components. These three modules determine a higher-level overall linear structure of signal propagation from the in-component (which includes the input CO_2_ signal) to the feedback-rich SCC, and finally to the out-component and closure (Figure 2B). The HCO_3_^−^ activation of SLAC1 described in [36] forms a unique edge that connects the CO_2_ in-component directly to the out-component.

### Construction of the dynamic model

A network of signaling elements reveals the interactions and regulatory relationships between them, yet this alone is not sufficient to capture the dynamic aspects of signal transduction. In the stomatal closure process the biological system represented by the network evolves from a state corresponding to open stomata (in the absence of a closure signal) to a state corresponding to closed stomata after sufficient time has elapsed in the presence of a closure signal. To capture the dynamic state transition of the CO_2_ network, we perform dynamic modeling, where we describe each node of the network with a state variable that represents its abundance or activity. In this way, the propagation of the high CO_2_ signal is captured in the model by a series of node state changes propagating through the network, eventually leading to stomatal closure. We use a Boolean model, i.e., we describe each node with binary states: ON or 1 for higher-than-threshold abundance or activity; OFF or 0 for lower-than-threshold abundance and activity. Despite its simplicity, Boolean dynamics has proved capable of capturing complex biological behaviors [37], e.g. cell differentiation as a consequence of multi-stability [38, 39], or sustained oscillations such as the cell cycle [40].

For each node, its state change is determined by the regulatory influences incident on it, which are captured by its Boolean regulatory function. The regulatory function uses the states of the node’s regulators in the network as inputs and combines these inputs using logical operators such that the function correctly reflects experimental observations. For example, the node “HCO_3_^−^” represents bicarbonate concentration in the guard cell. Its input nodes are “CO_2_ entry of cell” and “CA1/4” (the CO_2_-binding carbonic anhydrase enzymes CA1 and CA4). Since bicarbonate is produced from the reaction CO_2_ + H_2_O → HCO_3_^−^ + H^+^, catalyzed by CA1/4, we know that both CO_2_ entry into the cell and CA1/4 are required for the increase of intracellular bicarbonate concentration, thus the inputs are connected with a logical “AND” operator, and the regulatory function of node “HCO_3_^−^” is set as “*f_HCO_*_3-_ = CA1/4 AND CO2 entry of cell”. The comprehensive list of regulatory functions and their justifications are provided in Supporting Info S3. Each node’s regulatory function determines the future state of the node as determined by the current state of its regulators.

With the Boolean regulatory functions defined for each node, we can perform simulations of the signal transduction process over the entire CO_2_ network. We initialize the *in silico* stomata in a pre-stimulus, i.e. open state, with an ON state for closure-inhibiting nodes (namely ABI1, ABI2, CBC1/2, HAB1, HT1, and H^+^ ATPase) and other nodes OFF (see Supporting Info S3 for the complete initial condition). We deploy a discrete-time simulation, with stochastic random order asynchronous update, and simulate the network model for 30 timesteps (see Methods). Each simulated trajectory of the model will eventually converge into a stabilized behavior of the system as a whole, called an attractor. The most intuitive attractor is a steady state, where all nodes stabilize in a fixed state. Alternatively, a system can also have an oscillatory attractor, where a fraction of nodes indefinitely oscillate between 0 (OFF) and 1 (ON). A system can have multiple attractors (called multi-stability), which may be reachable from the same initial state if the system’s timing is stochastic. We obtain the consensus behavior of the system by performing 1000 replicate simulations, i.e., 1000 “*in silico* stomata” simulations starting in the same pre-stimulus state, then calculating the percentage of node activation at each timestep as our metric, meaning the percentage of simulations in which the node (e.g. Closure) is in state 1 (ON). In this way, oscillations and multi-stability with different node states will display a percentage of node activation with a value between 0% and 100%. We demonstrate simulated high CO_2_ induced stomatal closure and lack of closure under ambient CO_2_ in Figure 3 Figure 2B, and show a summary of node states in the corresponding attractors in Figure 3C.

**Figure 3.**
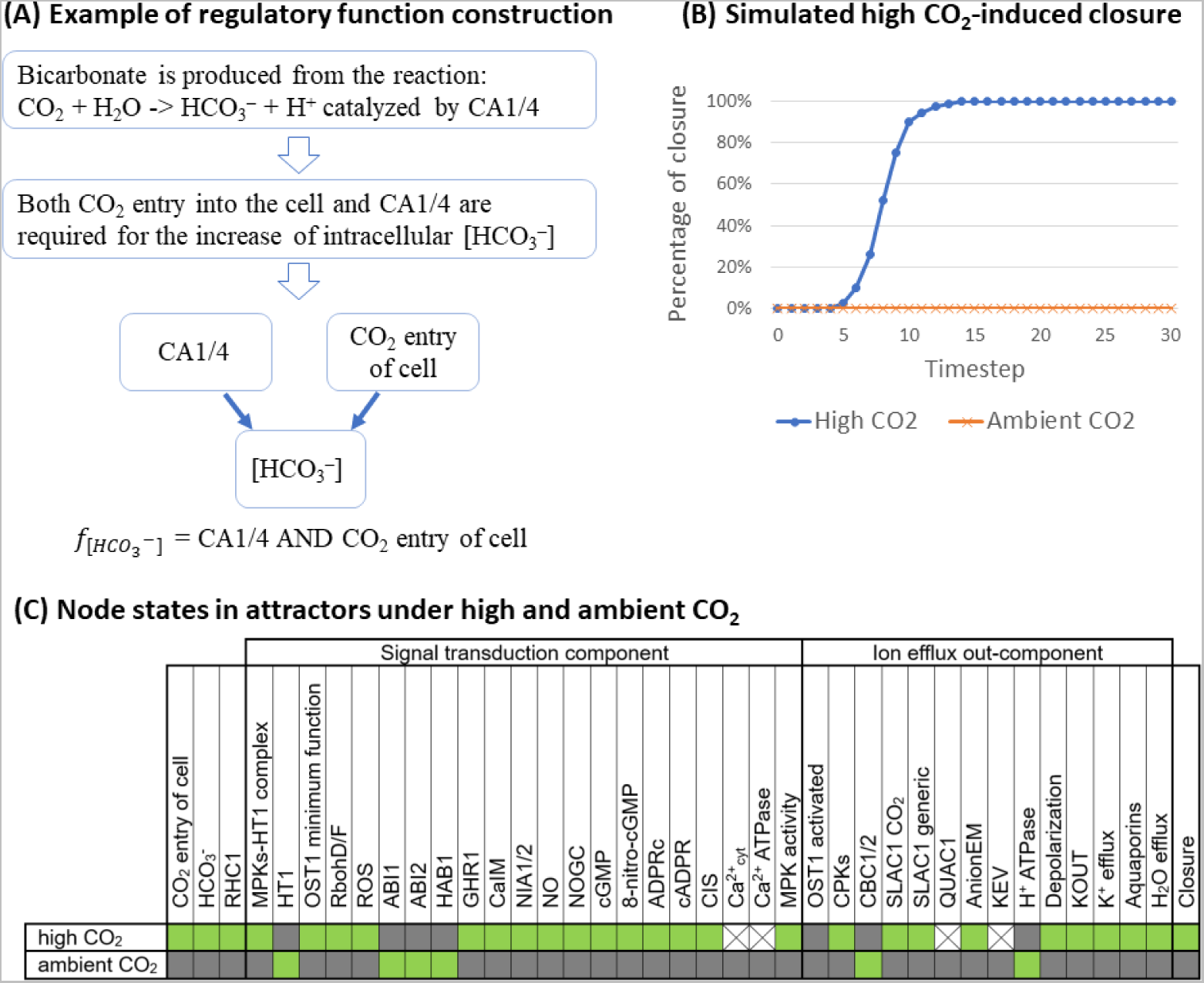
(A) Example demonstrating how a Boolean regulatory function is constructed from observations; (B) A demonstration of model simulation, capturing stomatal closure under high CO_2_, and lack of closure under ambient CO_2_; (C) Summary of node states in the attractor with closure ON under high CO_2_, and the attractor with closure OFF under ambient CO_2_, found from the model simulations. A green box represents the ON state, grey represents the OFF state, and X represents oscillation.

In the simulated wild-type system (Figure 3B), the percentage of closure in response to high CO_2_ increases over the timesteps from 0% to 100%. Despite the deliberate choice of stochasticity in the timing, all the simulations converge into the same attractor, in which the node Closure is ON. The row marked “high CO_2_” in Figure 3C shows the states of all nodes in this attractor, recapitulating the biological knowledge regarding the activity of nodes in stomata that closed in response to high CO_2_. For example, the node “OST1 activated” is OFF in this closure attractor, meaning that OST1 does not become activated under high CO_2_, consistent with recent publications [32, 33]. The node Ca^2+^_cyt_ oscillates between ON and OFF, capturing transient cytosolic Ca^2+^ concentration increases that have been observed during CO_2_ response [17, 18]. We observe that the node “SLAC1 generic” is activated (as all of its regulators, including CPKs, are indirectly activated by high CO_2_). Conversely, in the simulation curve corresponding to ambient CO_2_ (represented as the inactivity of the node “high CO_2_”) the percentage of closure remains 0% for the entire duration of the simulation, consistent with the expectation that stomata are open in the absence of CO_2_ elevation. The row marked “ambient CO_2_” in Figure 3C shows the states of all nodes in the attractor under ambient CO_2_.

### Validation of the network model

We assembled all experimental evidence related to high CO_2_ induced stomatal closure that was not used in model construction (see the entries with “V” for validation, in Supporting Info S1), and checked if the model simulations of the relevant settings yield results consistent with the experimental evidence. An important concept that can be insufficiently appreciated by experimentalists is that all the pathway-level effects (e.g., the effect of a node manipulation on high CO_2_ induced stomatal closure) are emergent properties of the system that arise from multiple interactions; thus, the fact that the regulatory functions of each node are informed by experimental evidence is not sufficient to guarantee consistency with pathway-level observations.

The summary of validation and consistency is presented in Table 2. In a total of 47 settings, there are 43 cases of consistency between the experimental and simulation results and 4 inconsistent cases, yielding an accuracy of 43/47 = 91%. The high validation accuracy shows that our model correctly captures most known signaling behaviors in CO_2_-induced stomatal closure. Note that a few comparisons could be consistent by construction if the observed regulator node directly regulates the target node in the model. When assessing the validity of our model, we exclude this type of trivial consistency. A more detailed validation table including the trivially consistent cases, a description of each experimental observation, and explanations are provided in Supporting Info S5.

**Table 2.**
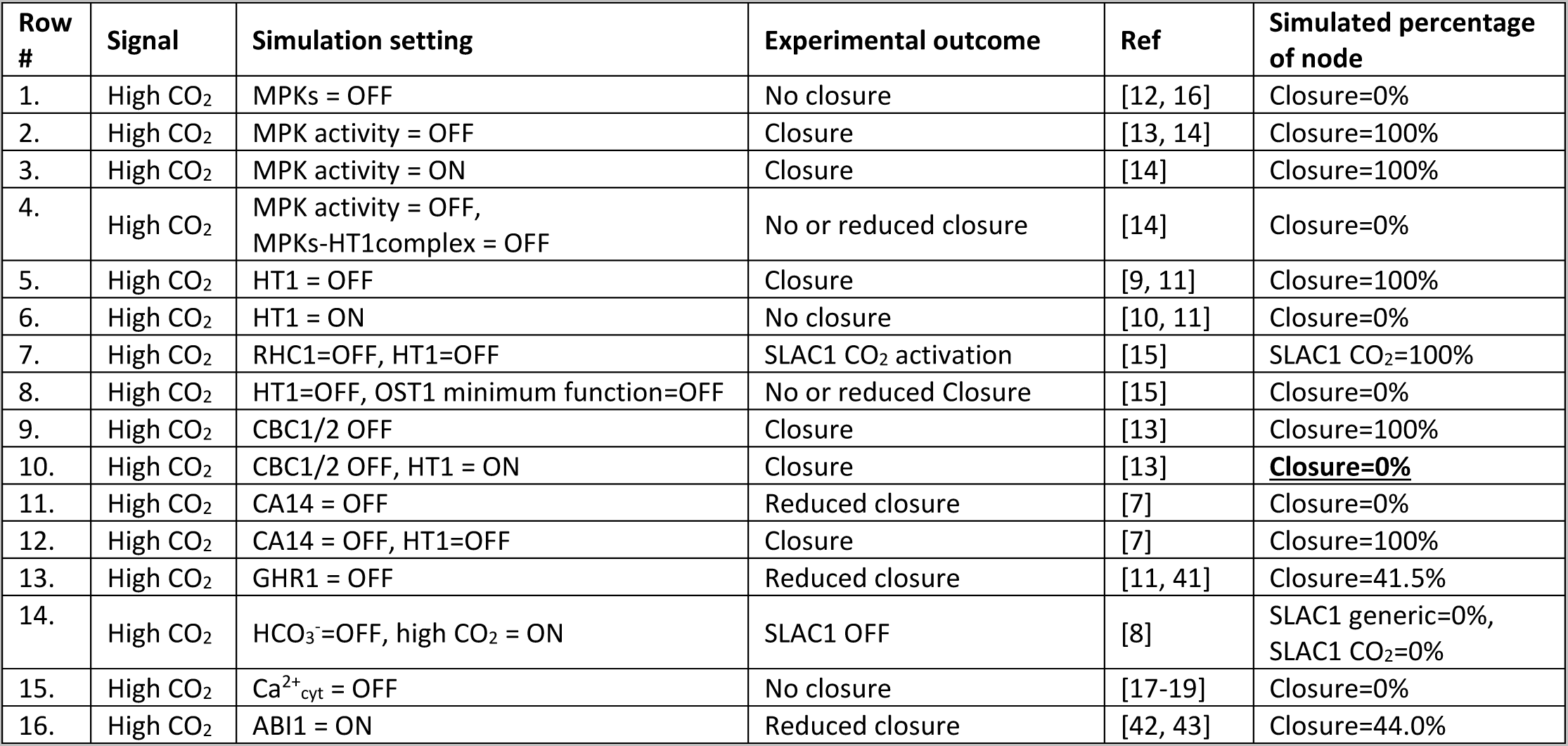
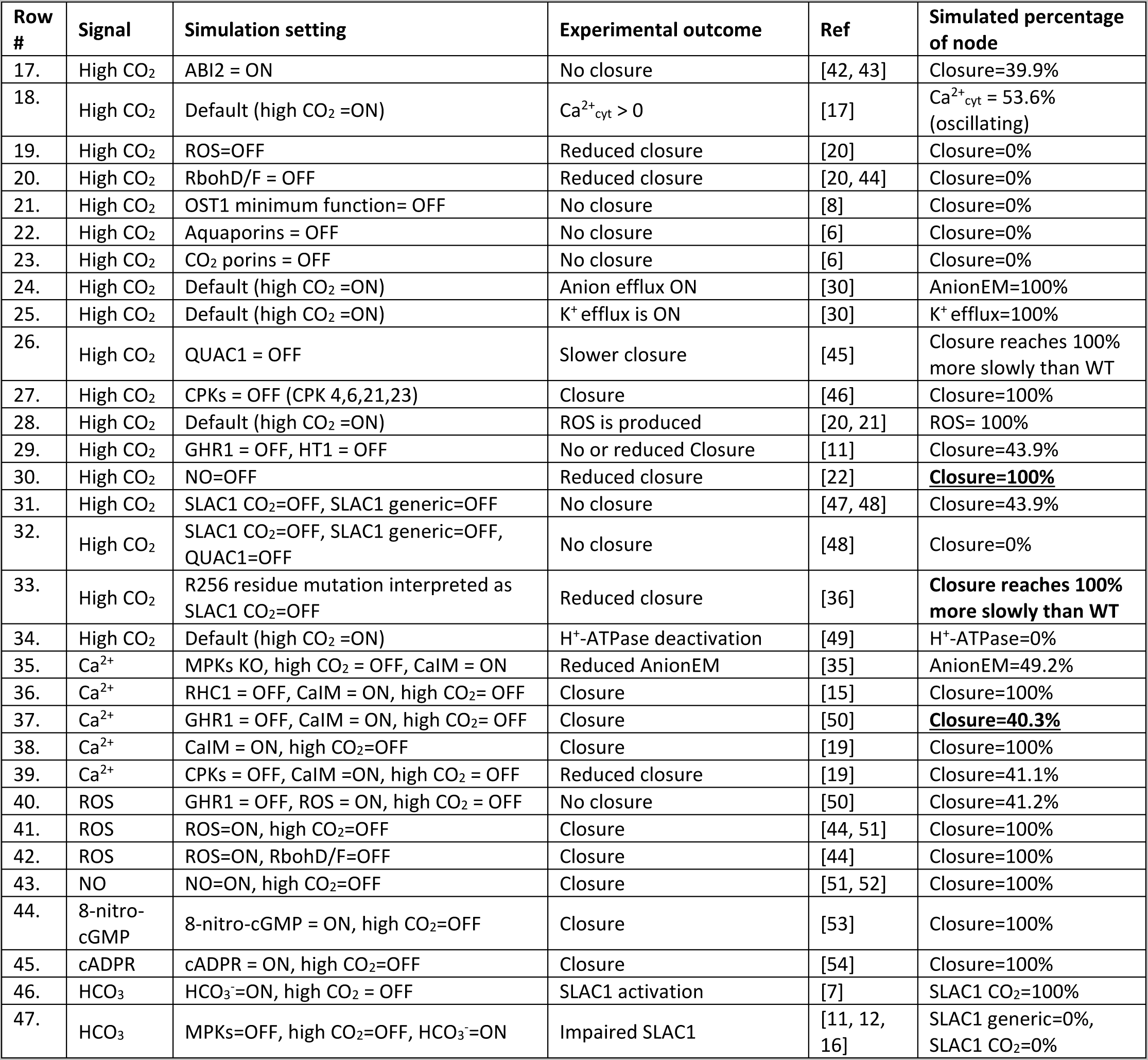
Comprehensive validation comparing simulations against known experimental observations. In the 47 total simulations in the table, 43 are consistent, 4 are inconsistent (marked with bold, underlined font), with accuracy 43/47 = 91% here.

| Row # | Signal | Simulation setting | Experimental outcome | Ref | Simulated percentage of node |
| --- | --- | --- | --- | --- | --- |
| 1. | High CO<sub>2</sub> | MPKs = OFF | No closure | [12, 16] | Closure=0% |
| 2. | High CO<sub>2</sub> | MPK activity = OFF | Closure | [13, 14] | Closure=100% |
| 3. | High CO<sub>2</sub> | MPK activity = ON | Closure | [14] | Closure=100% |
| 4. | High CO<sub>2</sub> | MPK activity = OFF, MPKs-HT1complex = OFF | No or reduced closure | [14] | Closure=0% |
| 5. | High CO<sub>2</sub> | HT1 = OFF | Closure | [9, 11] | Closure=100% |
| 6. | High CO<sub>2</sub> | HT1 = ON | No closure | [10, 11] | Closure=0% |
| 7. | High CO<sub>2</sub> | RHC1=OFF, HT1=OFF | SLAC1 CO<sub>2</sub> activation | [15] | SLAC1 CO<sub>2</sub>=100% |
| 8. | High CO<sub>2</sub> | HT1=OFF, OST1 minimum function=OFF | No or reduced Closure | [15] | Closure=0% |
| 9. | High CO<sub>2</sub> | CBC1/2 OFF | Closure | [13] | Closure=100% |
| 10. | High CO<sub>2</sub> | CBC1/2 OFF, HT1 = ON | Closure | [13] | **Closure=0%** |
| 11. | High CO<sub>2</sub> | CA14 = OFF | Reduced closure | [7] | Closure=0% |
| 12. | High CO<sub>2</sub> | CA14 = OFF, HT1=OFF | Closure | [7] | Closure=100% |
| 13. | High CO<sub>2</sub> | GHR1 = OFF | Reduced closure | [11, 41] | Closure=41.5% |
| 14. | High CO<sub>2</sub> | HCO<sub>3</sub><sup>-</sup>=OFF, high CO<sub>2</sub> = ON | SLAC1 OFF | [8] | SLAC1 generic=0%, SLAC1 CO<sub>2</sub>=0% |
| 15. | High CO<sub>2</sub> | Ca<sup>2+</sup><sub>cyt</sub> = OFF | No closure | [17-19] | Closure=0% |
| 16. | High CO<sub>2</sub> | ABI1 = ON | Reduced closure | [42, 43] | Closure=44.0% |
| 17. | High CO<sub>2</sub> | ABI2 = ON | No closure | [42, 43] | Closure=39.9% |
| 18. | High CO<sub>2</sub> | Default (high CO<sub>2</sub> =ON) | Ca<sup>2+</sup><sub>cyt</sub> > 0 | [17] | Ca<sup>2+</sup><sub>cyt</sub> = 53.6% (oscillating) |
| 19. | High CO<sub>2</sub> | ROS=OFF | Reduced closure | [20] | Closure=0% |
| 20. | High CO<sub>2</sub> | RbohD/F = OFF | Reduced closure | [20, 44] | Closure=0% |
| 21. | High CO<sub>2</sub> | OST1 minimum function= OFF | No closure | [8] | Closure=0% |
| 22. | High CO<sub>2</sub> | Aquaporins = OFF | No closure | [6] | Closure=0% |
| 23. | High CO<sub>2</sub> | CO<sub>2</sub> porins = OFF | No closure | [6] | Closure=0% |
| 24. | High CO<sub>2</sub> | Default (high CO<sub>2</sub> =ON) | Anion efflux ON | [30] | AnionEM=100% |
| 25. | High CO<sub>2</sub> | Default (high CO<sub>2</sub> =ON) | K<sup>+</sup> efflux is ON | [30] | K<sup>+</sup> efflux=100% |
| 26. | High CO<sub>2</sub> | QUAC1 = OFF | Slower closure | [45] | Closure reaches 100% more slowly than WT |
| 27. | High CO<sub>2</sub> | CPKs = OFF (CPK 4,6,21,23) | Closure | [46] | Closure=100% |
| 28. | High CO<sub>2</sub> | Default (high CO<sub>2</sub> =ON) | ROS is produced | [20, 21] | ROS= 100% |
| 29. | High CO<sub>2</sub> | GHR1 = OFF, HT1 = OFF | No or reduced Closure | [11] | Closure=43.9% |
| 30. | High CO<sub>2</sub> | NO=OFF | Reduced closure | [22] | **Closure=100%** |
| 31. | High CO<sub>2</sub> | SLAC1 CO<sub>2</sub>=OFF, SLAC1 generic=OFF | No closure | [47, 48] | Closure=43.9% |
| 32. | High CO<sub>2</sub> | SLAC1 CO<sub>2</sub>=OFF, SLAC1 generic=OFF, QUAC1=OFF | No closure | [48] | Closure=0% |
| 33. | High CO<sub>2</sub> | R256 residue mutation interpreted as SLAC1 CO<sub>2</sub>=OFF | Reduced closure | [36] | **Closure reaches 100% more slowly than WT** |
| 34. | High CO<sub>2</sub> | Default (high CO<sub>2</sub> =ON) | H<sup>+</sup>-ATPase deactivation | [49] | H<sup>+</sup>-ATPase=0% |
| 35. | Ca<sup>2+</sup> | MPKs KO, high CO<sub>2</sub> = OFF, CaIM = ON | Reduced AnionEM | [35] | AnionEM=49.2% |
| 36. | Ca<sup>2+</sup> | RHC1 = OFF, CaIM = ON, high CO<sub>2</sub>= OFF | Closure | [15] | Closure=100% |
| 37. | Ca<sup>2+</sup> | GHR1 = OFF, CaIM = ON, high CO<sub>2</sub>= OFF | Closure | [50] | **Closure=40.3%** |
| 38. | Ca<sup>2+</sup> | CaIM = ON, high CO<sub>2</sub>=OFF | Closure | [19] | Closure=100% |
| 39. | Ca<sup>2+</sup> | CPKs = OFF, CaIM =ON, high CO<sub>2</sub> = OFF | Reduced closure | [19] | Closure=41.1% |
| 40. | ROS | GHR1 = OFF, ROS = ON, high CO<sub>2</sub> = OFF | No closure | [50] | Closure=41.2% |
| 41. | ROS | ROS=ON, high CO<sub>2</sub>=OFF | Closure | [44, 51] | Closure=100% |
| 42. | ROS | ROS=ON, RbohD/F=OFF | Closure | [44] | Closure=100% |
| 43. | NO | NO=ON, high CO<sub>2</sub>=OFF | Closure | [51, 52] | Closure=100% |
| 44. | 8-nitro-cGMP | 8-nitro-cGMP = ON, high CO<sub>2</sub>=OFF | Closure | [53] | Closure=100% |
| 45. | cADPR | cADPR = ON, high CO<sub>2</sub>=OFF | Closure | [54] | Closure=100% |
| 46. | HCO<sub>3</sub> | HCO<sub>3</sub>=ON, high CO<sub>2</sub> = OFF | SLAC1 activation | [7] | SLAC1 CO<sub>2</sub>=100% |
| 47. | HCO<sub>3</sub> | MPKs=OFF, high CO<sub>2</sub>=OFF, HCO<sub>3</sub>=ON | Impaired SLAC1 | [11, 12, 16] | SLAC1 generic=0%, SLAC1 CO<sub>2</sub>=0% |

### Network-based analysis identifies the dynamic repertoire and decision-making mechanisms of the CO_2_ signaling system

In the previous section we showed that our model can accurately reproduce existing experimental observations. Next, we explore the explanatory power of the model and its predictions about the biological system’s response in conditions for which experimental observations are not yet available.

We start by determining the full attractor repertoire of the model. The attractor repertoire of the model is important because attractors correspond to biological phenotypes, e.g., open or closed stomata are represented by two different attractors in the CO_2_ network model. Finding the attractor repertoire is the first step toward understanding how a biological system makes decisions on how to switch between different phenotypes. Understanding the decision-making behind attractor transitions of a biological system can reveal the key signal transduction mechanisms and suggest potential interventions to guide the system into a more desirable state. Finding attractors by simulations is difficult because the state space of a network model increases exponentially with its size, e.g., the CO_2_ network model with 48 nodes has a total of 2^48^ states. An alternative and efficient way to find the attractors of a system is to utilize the network topology via stable motif analysis, developed by the Albert group [55]. Stable motifs are special feedback loops in a dynamic model that can self-sustain once their constituent nodes attain an associated state (see Methods). The important feature of stable motifs is that they are irreversible: after locking in they become independent of the rest of the system. This feature implies that the lock-in of a stable motif is like a decision of the system, and a series of stable motif lock-ins will lead to an attractor [26, 28, 55]. Therefore, finding all stable motifs and their lock-in sequence allows comprehensive identification of attractors.

### Stable motifs identify the dynamic repertoire and decision-making of the system

Whether in ambient or high CO_2_, the unperturbed (wild type) system has a stable motif (see Figure 4A) that includes the activity of RbohD/F and ROS, the minimum functional level of OST1, and the inactivity of the three PP2C protein phosphatases (ABI1, ABI2 and HAB1). We will refer to this stable motif as the Main Stable Motif (MSM). The lock-in of the MSM is observed in the closure ON attractor (the first row of Figure 3C). The opposite states of the nodes involved in the MSM, namely the inactivity of RbohD/F, ROS, CaIM, and GHR1, the lack of OST1 function, and the activation of one of the PP2C protein phosphatases, form three instances of a weaker type of stable motif, called a conditionally stable motif [56] (see Methods). Figure 4B indicates the superposition of the three conditionally stable motifs and shows that they can lock-in if HT1 is ON. The lock-in of these conditionally stable motifs leads to the attractor with closure OFF (second row in Fig. 3C), which is consistent with the CO_2_-independent high stomatal conductance of the *ht1-3D* and *ht1-8D* dominant mutants [10, 11]. Importantly, the MSM and the three conditionally stable motifs are mutually exclusive. Since the lock-in of a stable motif is irreversible, the choice between them constitutes the decision-making process of the CO_2_ signaling network (Figure 4C). The system has additional (conditionally) stable motifs that contribute in specific circumstances; these are described in section II of Supporting Info S4.

**Figure 4.**
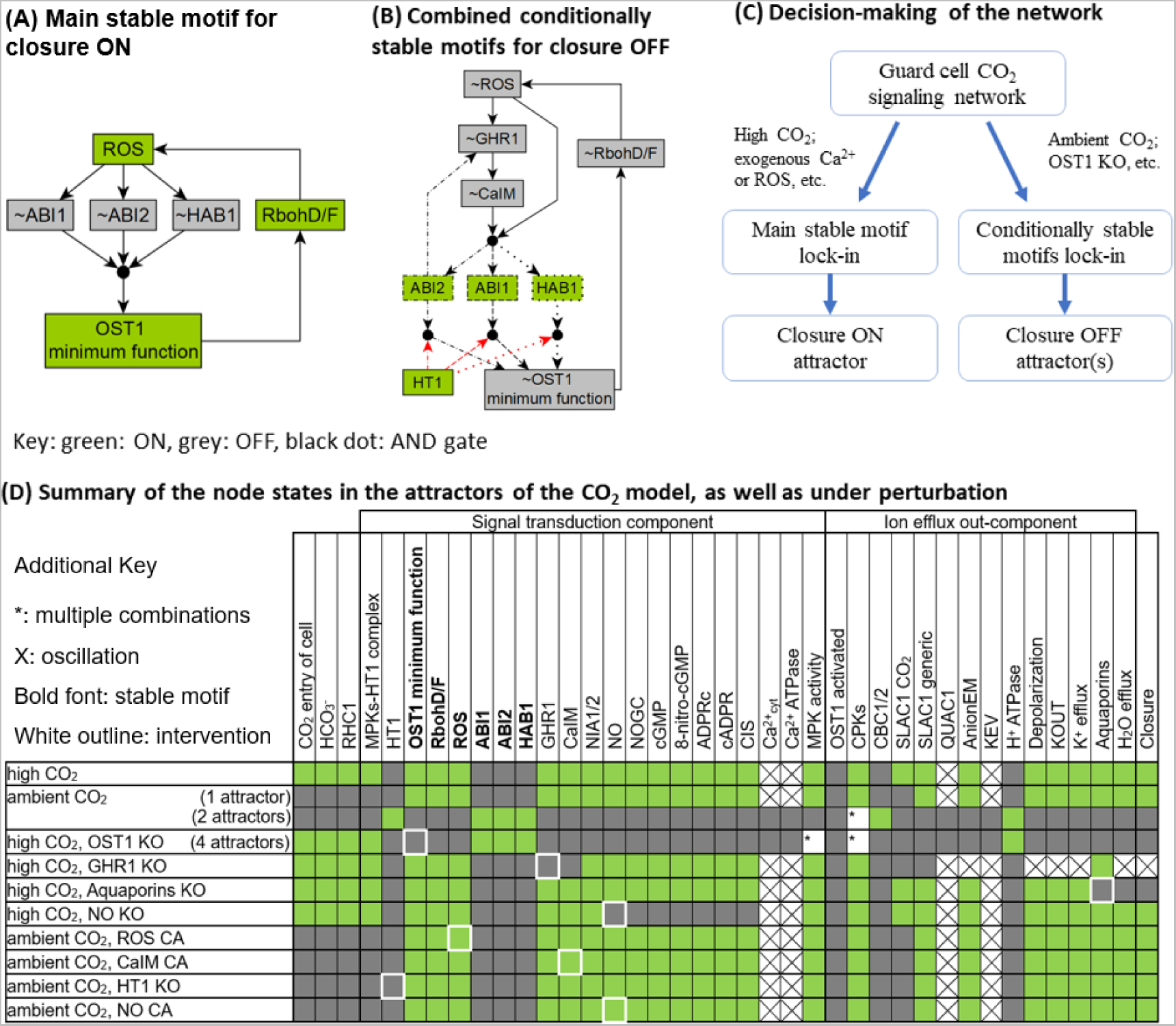
Stable motifs of the network model. (A) The main stable motif that leads to closure in the dynamic model. (B) Superposition of the three overlapping conditionally stable motifs that lead to attractors that represent open *in silico* stomata. Each conditionally stable motif contains a different PP2C protein family member, indicated by dashed, dash-dotted or dotted outline and edges, respectively. In the stable motifs, nodes whose name is preceded by ~ are inactive in the corresponding motif. The black circles indicate that the regulatory effects incident on a node form an AND gate; for example, ABI1 will activate only if ROS and CaIM are simultaneously inactive. The red edges starting from HT1 indicate that HT1=ON is the condition necessary for the conditionally stable motifs. (C) Decision-making diagram of the CO_2_ signaling network. (D) Summary of the node states in the attractors of the system. Entries that contain a white box with * summarize multiple attractors in which the marked nodes take either on or off states.

Identifying the stable motifs allows comprehensive identification of all attractors of the system, using the algorithm developed in [27]. We first confirm the unique Closure ON attractor in the presence of high CO_2_ (Figure 4D), where high CO_2_ leads to the locking-in of the Main Stable Motif and precludes the activation of the three conditionally stable motifs. The MSM leads to the activation of anion flow through the SLAC1 channel and to K^+^ efflux, as well as to the activation of Aquaporins, which together satisfy all requirements to activate the node Closure. In the case of ambient CO_2_, stable motif analysis identifies 3 attractors. Two of them have the node Closure OFF, corresponding to the absence of stomatal closure observed under ambient CO_2_. These two attractors differ only in the state of CPKs (either ON or OFF). Interestingly, the 3^rd^ attractor under ambient CO_2_ has the node Closure ON, and is highly similar to the closure ON attractor under high CO_2_ --they differ only in the CO_2_ in-component (see Figure 4D, row 2 vs. row 1). Trajectories starting from the pre-stimulus initial state used in our simulation cannot reach this attractor under ambient CO_2_, consistent with biological reality and with the attractor found by simulation (Figure 3C). Yet, a different initial state can lead to this attractor (see Supporting Info S3). An intervention (e.g., sustained activation of a node) can also induce stomatal closure under ambient CO_2_, as we will present later.

### Logic diagrams elucidate signal transduction mechanisms and the roles of driver nodes

To better understand how the system reaches its attractors (e.g., the closure ON attractor under ambient CO_2_), we introduce logic diagrams illustrating the signaling mechanisms (Figure 5). A logic diagram (see Methods) is an integration of the network and node states that yield a specific outcome (e.g., the ON state of the node Closure), constructed based on the framework of the “expanded network” developed by the Albert group [55, 57, 58]. Edges in the logic diagrams are logical implications (i.e., sufficient activations), and black dots represent AND gates. To make the logic diagrams easier to parse, we also applied network reduction techniques that do not change the dynamic behavior of the system [59], and we merged nodes that behave similarly or as a group (e.g. PP2Cs, see Methods).

**Figure 5.**
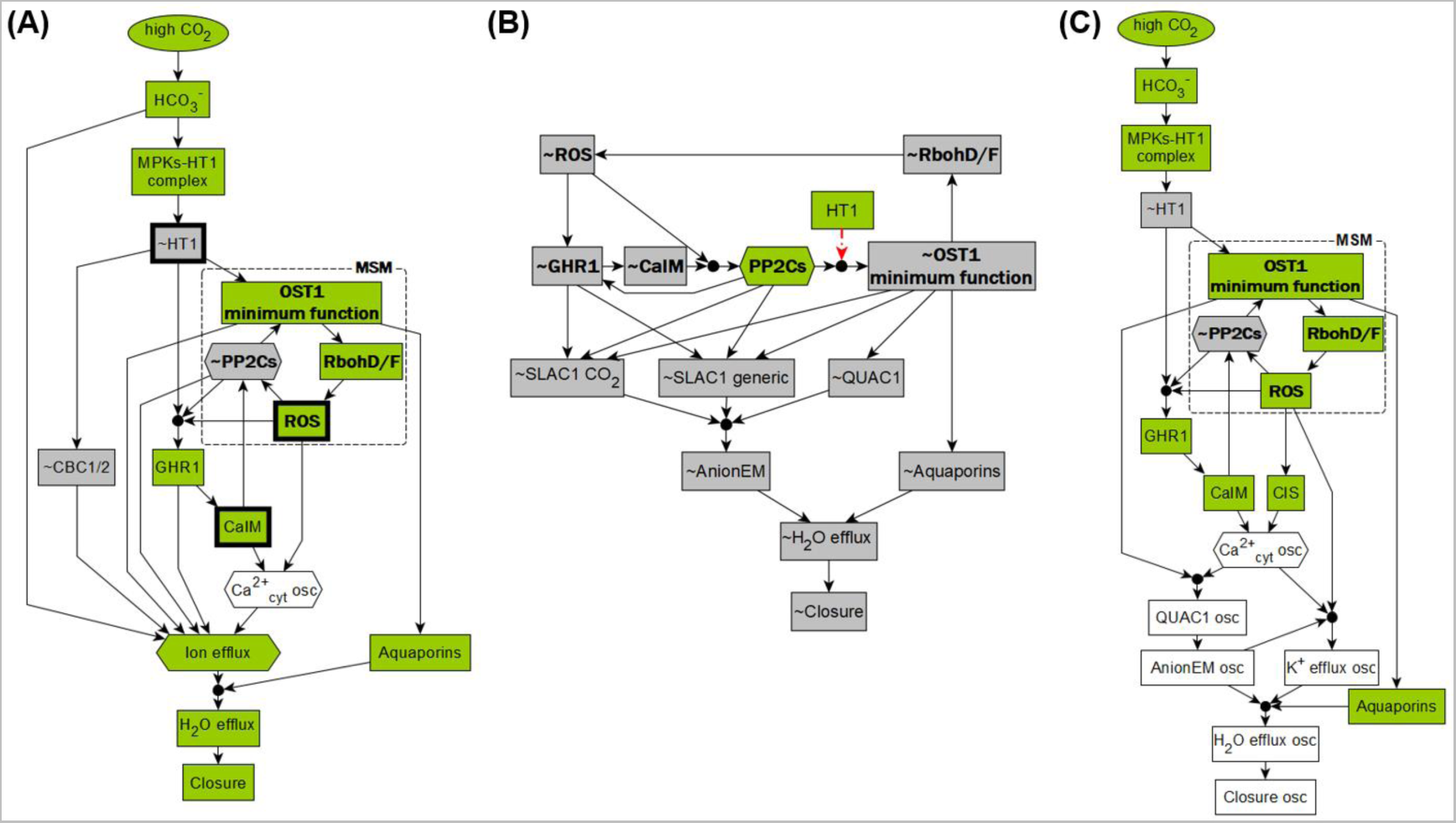
The logic diagrams show the mechanisms that lead to various stomatal behaviors. (A) Logic diagram of high CO_2_-induced stomatal closure. It also indicates that a closure attractor can be reached under ambient CO_2_ if the Main Stable Motif is activated by an internal or external driver. Three such drivers (out of 17) are indicated by bold outlines. (B) Diagram of the mechanism for the lack of closure due to the activation of the conditionally stable motifs, e.g., under ambient CO_2_ or for *ost1* KO. (C) Logic diagram of the predicted mechanism of unstable closure that would arise with inactivity of the SLAC1 channels. Green nodes indicate the active state and grey means the inactive state; white nodes oscillate.

As shown in Figure 5A, high CO_2_ activates the MSM (shown with bold font), which leads to ion and water efflux, and thus to an attractor with the node Closure ON. Figure 5B shows the mechanism for the lack of stomatal closure, e.g., under ambient CO_2_. When HT1 is ON, the conditionally stable motifs (bold-font nodes, with their dependence on HT1=ON shown with red dotted edges) can lock in and cause the inactivity of the SLAC1 and QUAC1 channels, thus reaching a Closure OFF attractor. The knockout of OST1 also activates the conditionally stable motifs and leads to a similar Closure OFF attractor (see the third entry of Figure 4D). In addition to attractors with constant closure values (constant ON or OFF), the system also allows attractors with an unstable closure value that oscillates between ON and OFF indefinitely. Figure 5C shows the logic diagram of the predicted mechanism of unstable Closure state that could arise from the sustained inactivity of the SLAC1 channels (due, e.g., to knockout of GHR1, see the fourth entry of Figure 4D). The oscillation of Ca^2+^_cyt_ leads to oscillating QUAC1 and K^+^ efflux, which in turn lead to an oscillating Closure state. Had SLAC1 been active, it would have sustained anion efflux, leading to sustained Closure ON.

In addition to finding attractors, stable motif analysis can also pinpoint key nodes called drivers, which determine the decisions in the system: once a driver node is sustained in a certain state, it can lock in a corresponding stable motif, forcing a decision in the system. Thus, external control of driver nodes can drive the system from one attractor to another [25, 27]. For example, ROS (marked in bold outline) is an internal driver of the MSM, as it inhibits all three PP2C proteins, which then allows OST1 activity, which in turn activates RbohD/F, closing the cycle of self-sustained activity. Furthermore, CaIM (Ca^2+^ influx through the membrane) is an external driver of the MSM, as it inhibits the PP2Cs. When ROS or Ca^2+^ is externally provided, it will lock in the MSM, thus lead to a closure ON attractor, even in ambient CO_2_ (see Figure 4D). This explains the experimentally observed ROS or Ca^2+^ induced stomatal closure under ambient CO_2_ [19, 44, 60]. Moreover, ~HT1 (the OFF state of HT1) is also an external driver of the MSM, and leads to the same closure ON attractor as providing ROS or Ca^2+^ (see Figure 4D), which explains the low stomatal conductance observed in the *ht1-2* kinase dead mutant regardless of the CO_2_ concentration [9].

### Our model predicts how the perturbation of signaling elements affects high CO_2_ response

A model is most helpful in predicting unknown behaviors of a system. We first demonstrate the predictive power of our model by simulating the guard cell response to high CO_2_ under systematic single node perturbations. These simulations reflect how mutations or interventions could affect high CO_2_ induced closure. We simulate all single interventions that abolish a node’s expression or activity as maintaining the node in the state 0 (OFF) and denote it “KO”; we also simulate all single interventions that make a node constitutively active by maintaining the node in the state 1 (ON) and denote it “CA”. To quantify the stomatal closure response, we develop an improved approach from [3] combining three metrics: the final closure state, the cumulative percentage of closure (CPC), as well as the attractor repertoire of the system (see Methods). The CPC value represents the fraction of simulations in which Closure = 1, summed over 30 timesteps, thus it ranges from 0 to 30. As before [3, 61], we observe and categorize four typical behaviors different than the wild type (WT) response: (1) we call the behavior in which the percentage of closure increases faster than in the WT system “hypersensitivity”, and (2) the behavior in which the percentage of closure increases more slowly than in the WT system “hyposensitivity”; (3) we refer to the behavior in which the percentage of closure converges to or oscillates around an intermediate value “reduced closure”; (4) and finally the category “no closure” is the lack of closure in simulated stomata. We present the summarized high CO_2_ closure response categories under systematic perturbations in Figure 6A. The complete perturbation results can be found in Supporting Info S6. We illustrate simulated response curves of these categories in Figure 6B: hypersensitivity under external Ca^2+^ (CaIM CA), hyposensitivity under CaIM KO, reduced closure under H^+^-ATPase KO, and no closure under MPKs KO (*mpk4/12* mutant). These perturbation response categories offer a way to characterize each signaling element’s contribution to high CO_2_ induced closure, from weakest (nodes whose KO and CA are in the “Close to WT” category, such as cGMP) to strongest (nodes whose KO is in the “No closure” category and whose CA is in the “Hypersensitivity” category, such as ROS).

**Figure 6.**
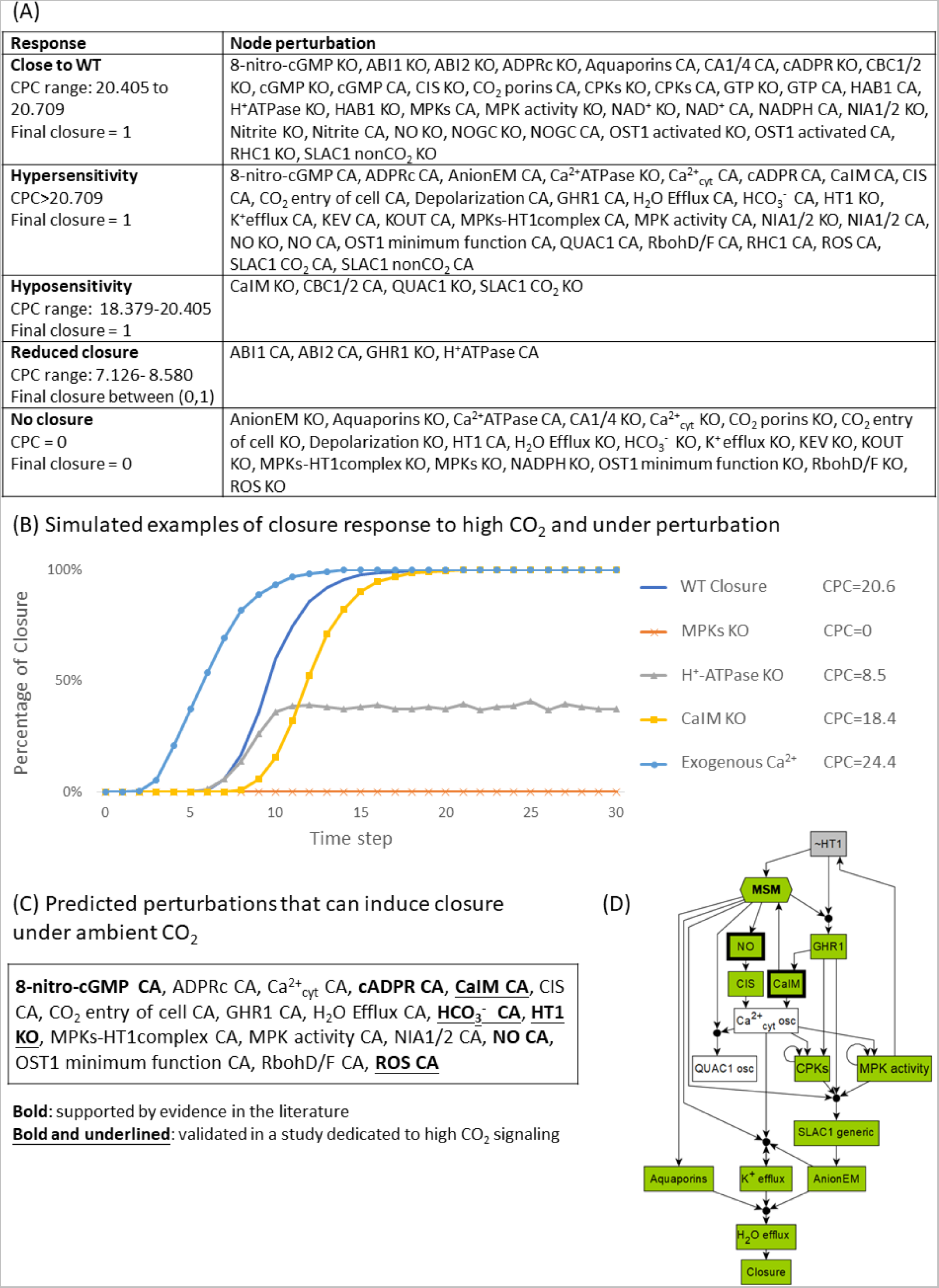
Systematic perturbation under high CO_2_ with illustrations. (A) Response categorization for systematic perturbations of all nodes (both KO and CA perturbations). (B) Simulation curves under high CO_2_ that exemplify response types: normal WT closure, hypersensitivity under external Ca^2+^ (Ca^2+^ influx across the membrane (CaIM) CA), hyposensitivity under CaIM KO, reduced CO_2_ sensitivity for H^+^-ATPase, or insensitivity to CO_2_ under MPKs KO. (C) Model simulation predicts 18 interventions that could induce stomatal closure under ambient CO_2_. Bolded interventions have supporting evidence in previous publications, and bolded plus underlined interventions have been validated to induce closure under ambient CO_2_. (D) Logic diagram of the mechanism of closure in ambient CO_2_. The node “MSM” is a merged representation of the Main Stable Motif. Bold outlines indicate 2 of the 18 interventions that can simultaneously activate the MSM and inactivate HT1 under ambient CO_2_. For example, NO leads to oscillations in Ca^2+^, which lead to MPK activity, which inactivates HT1, which then activates the MSM and leads to closure.

### The five response categories are well-explained by our stable motif and network analysis

We find that in both hyper- and hypo-sensitivity cases, the final attractor of the system is almost the same as that of the wildtype system under high CO_2_; the final attractor only differs in the state of the perturbed node and up to a few other nodes regulated by it. As illustrated in the logic diagrams (Figure 5) earlier, the typical reason for hyper-sensitivity (i.e., faster closure) is the immediate accomplishment of a node state that would naturally happen only after a previous event; e.g., CaIM normally would only happen after GHR1 activation (Figure 5A), but it happens immediately with CaIM CA. The mechanism for slower closure is the elimination of one of two redundant signaling pathways: for example, CaIM KO will eliminate one pathway to increase the cytosolic Ca^2+^ level (a necessary condition of closure); the alternative pathway to activate Ca^2+^ is by CIS (Ca^2+^ influx from stores). The model prediction that perturbation of specific anion channels, e.g. QUAC1 KO or R256 residue KO (interpreted as SLAC1 CO_2_=OFF) leads to hyposensitivity is due to the same type of mechanism. Reduced closure occurs when the state of the node Closure oscillates in all attractors, such as for GHR1 KO (see row 6 in Figure 4D and Figure 5C). GHR1 KO leads to the deactivation of the SLAC1 anion channel, leaving QUAC1 as the only conduit of anion flow, and the activity of QUAC1 follows the Ca^2+^ oscillation, resulting in an attractor with oscillatory closure. The final category of behavior, “no closure”, is the lack of closure in all simulated stomata; the system converges into an attractor (or mix of attractors) with closure OFF. The mechanism that leads to this outcome is disruption of ion or water efflux, either through a mutation in a transporter (e.g., Aquaporins KO in Figure 4D), or indirectly through activation of the non-closure conditionally stable motifs (e.g., OST1 KO in Figure 4D; the mechanism is illustrated in Figure 5B).

### Our model predicts interventions that yield closure even under ambient CO_2_

We performed systematic perturbation simulations of the model under ambient CO_2_ (i.e. with the node “high CO_2_” set to OFF), and found 18 interventions that can result in Closure=ON in the simulations (Figure 6C; comprehensive results are in Supporting Info S6). Our stable motif analysis identifies that the reason for the effectiveness of these interventions is the activation of the MSM coupled with the inactivation of HT1 (see Figure 5A). HT1 KO itself is an example of such a stable-motif-activating intervention; this result agrees with the closed stomata or low stomatal conductance observed in the *ht1-2* mutant [9, 11]. Thus, the model predicts that constitutive activity of the nodes that lead (directly or indirectly) to the inhibition of HT1 can also drive the stable motifs and lead the system to closure in ambient CO_2_. Indeed, all the interventions listed in Figure 6C have this property. Figure 4D illustrates the Closure ON attractor that results from providing NO under ambient CO_2_ and Figure 6D illustrates the mechanism through which providing NO can lead to closure. In agreement with the model predictions, it was observed experimentally that provision of ROS [44] or CaIM (Ca^2+^ influx through the membrane, which is equivalent to providing external Ca^2+^) [19] results in closure under ambient CO_2_. The predicted closure-inducing effect of additional interventions is supported by experimental evidence, e.g. NO [51, 52], cADPR [54], 8-nitro-cGMP [53] were shown to induce closure as a signaling element of the ABA-induced closure network.

### Experimental validation of model predictions

Our model predicts that the activation of the MSM is an important part of the process of stomatal closure both in ambient and high CO_2_ (see Figure 5A and Figure 6D). A follow-up conclusion from this prediction is that earlier activation of the MSM would lead to earlier closure. One of the nodes whose constitutive activity is predicted to activate the MSM and lead to closure in ambient CO_2_ is NO. In our model, applying NO under high CO_2_ activates the MSM sooner. Hence, supplying NO is predicted to cause hypersensitivity of closure under high CO_2_ (i.e., faster high CO_2_-induced closure). To assess the validity of this prediction, we experimentally tested the effect of NO on high CO_2_ response by applying SNP (sodium nitroprusside dihydrate), an NO donor (Figure 7A, see Methods for experimental assay description). To observe any speed-up of the closure process, we used short treatments (10 and 20 minutes). We found that under high (800 ppm) CO_2_, the stomatal aperture under SNP treatment (orange and grey bars) is significantly lower than that without SNP treatment (blue bars, p=3.0e-2, Student’s t-test). Furthermore, the mean aperture corresponding to the 10 min combined treatment of 0.15 mM SNP and high CO_2_ is lower than the mean aperture corresponding to the 20 min high CO_2_ treatment (rightmost blue bar, p=4.3e-2, Student’s t-test), indicating that SNP treatment speeds up the high CO_2_ induced closure process, consistent with our model prediction.

**Figure 7.**
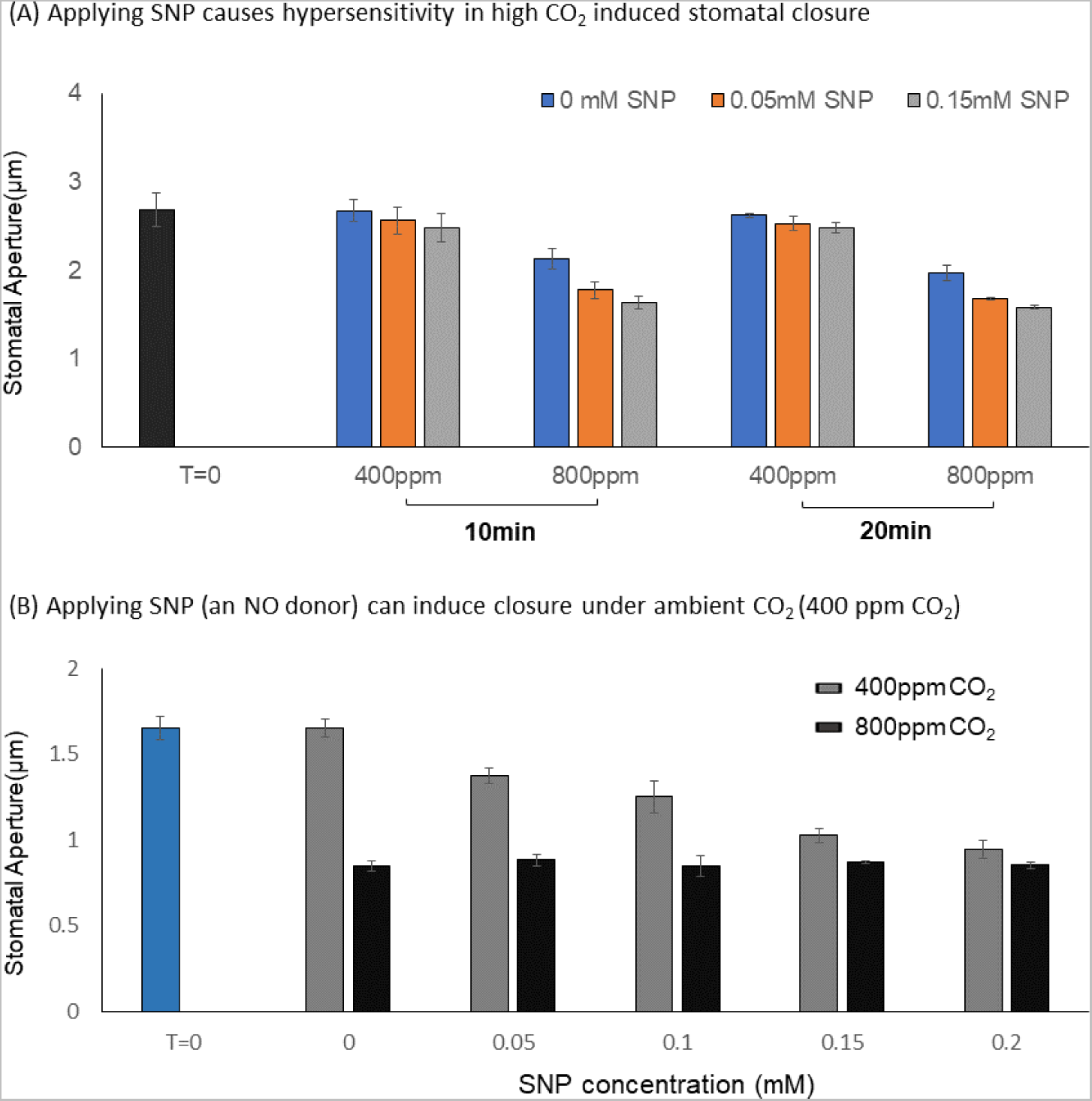
Experimental validation of the predicted closure responses when applying NO (via the NO donor sodium nitroprusside dihydrate, SNP) in ambient or high CO_2_. (A) As predicted by the model, applying SNP causes hypersensitivity in high CO_2_ induced stomatal closure: the combined treatment (orange and grey bars) leads to a lower mean aperture than high CO_2_ alone (blue bars), and a 10 min combined treatment leads to a lower mean aperture than a 20 min treatment of high CO_2_ alone. (B) Applying SNP (NO donor) under ambient (400 ppm) CO_2_ induces closure in a dose-dependent manner (grey bars), as predicted. The highest concentration of SNP yields a similar degree of closure as high (800 ppm) CO_2_ (black bars). The bars indicate the mean ± SE aperture size from at least three independent experiments with 30-40 apertures measured per treatment.

Furthermore, we experimentally validate the prediction that NO can induce closure under ambient CO_2_. Our model predicts that under ambient CO_2_, NO can still activate the MSM, consequently inducing closure (Figure 4C, Figure 6D). Previously, NO has been shown to induce closure as a signaling element of the ABA induced closure network [51, 52]. Yet, it has been unknown whether the degree of NO induced closure (aperture reduction) is comparable to high CO_2_ induced closure. We found that applying the NO donor SNP under ambient CO_2_ can indeed induce stomatal closure (Figure 7B, grey bars), as predicted by our model. The reduction in aperture caused by 0.2 mM SNP is statistically significant (p=6.9e-4, Student’s t-test), and close to the reduction in aperture caused by CO_2_ of 800 ppm (black bars). This result confirms the model prediction that NO can induce closure in ambient CO_2_.

Taken together, these experiments not only validate our model predictions regarding the effects of NO, but also support the model-predicted importance of the main stable motif.

### Stable motif analysis predicts a new regulatory relationship between NO and ABI2

Analysis of the inconsistencies between the model’s results and experimental observations can identify edges that are missing from the network model. The node NO offers an example of this kind as well. For example, while our model recapitulates NO-induced closure in ambient CO_2_, it does not capture the observation that NO depletion leads to loss of stomatal closure under elevated CO_2_ [22]. This discrepancy suggests that there is a not-yet-known regulatory effect of NO that makes it necessary for a closure mechanism. This unidentified regulatory effect of NO can be predicted from our stable motif analysis. In our current model under high CO_2_ and NO KO, the Main Stable Motif can still lock-in by the activation of its driver, the inactivity of HT1, thus leading to a closure ON attractor (Figure 8A, Figure 4D). According to current knowledge, NO participates in the pathway that leads to calcium influx from intracellular stores into the cytosol (CIS) [62]. Disruption of this pathway eliminates one mechanism of increasing the cytosolic Ca^2+^ level, but it does not perturb the Main Stable Motif. To be able to drive the system away from a closure ON attractor, NO KO should prevent the lock-in of the Main Stable Motif from activating closure (Figure 8B). Thus, we hypothesize that NO regulates at least one of the nodes in the MSM, namely, it activates ROS, RbohD/F, or OST1 minimum function, or inhibits ABI1, ABI2, or HAB1. Indeed, we experimentally found that NO inhibits ABI2’s phosphatase activity (Figure 8C, and see Methods). This result suggests that the mechanism (or one of multiple contributing mechanisms) leading to impairment of stomatal closure under NO knockout could be the increase of ABI2’s activity, which obstructs the minimal functional OST1 activity and precludes the lock-in of the MSM. As a decrease of ABI2’s phosphatase activity makes the activation of the MSM easier, this newly-discovered inhibitory regulation likely contributes to NO-induced stomatal closure as well. The experimental confirmation of this stable-motif-based model prediction highlights the power of our model in predicting/prioritizing potential regulatory relationships and understanding the mechanisms of signal transduction.

**Figure 8.**
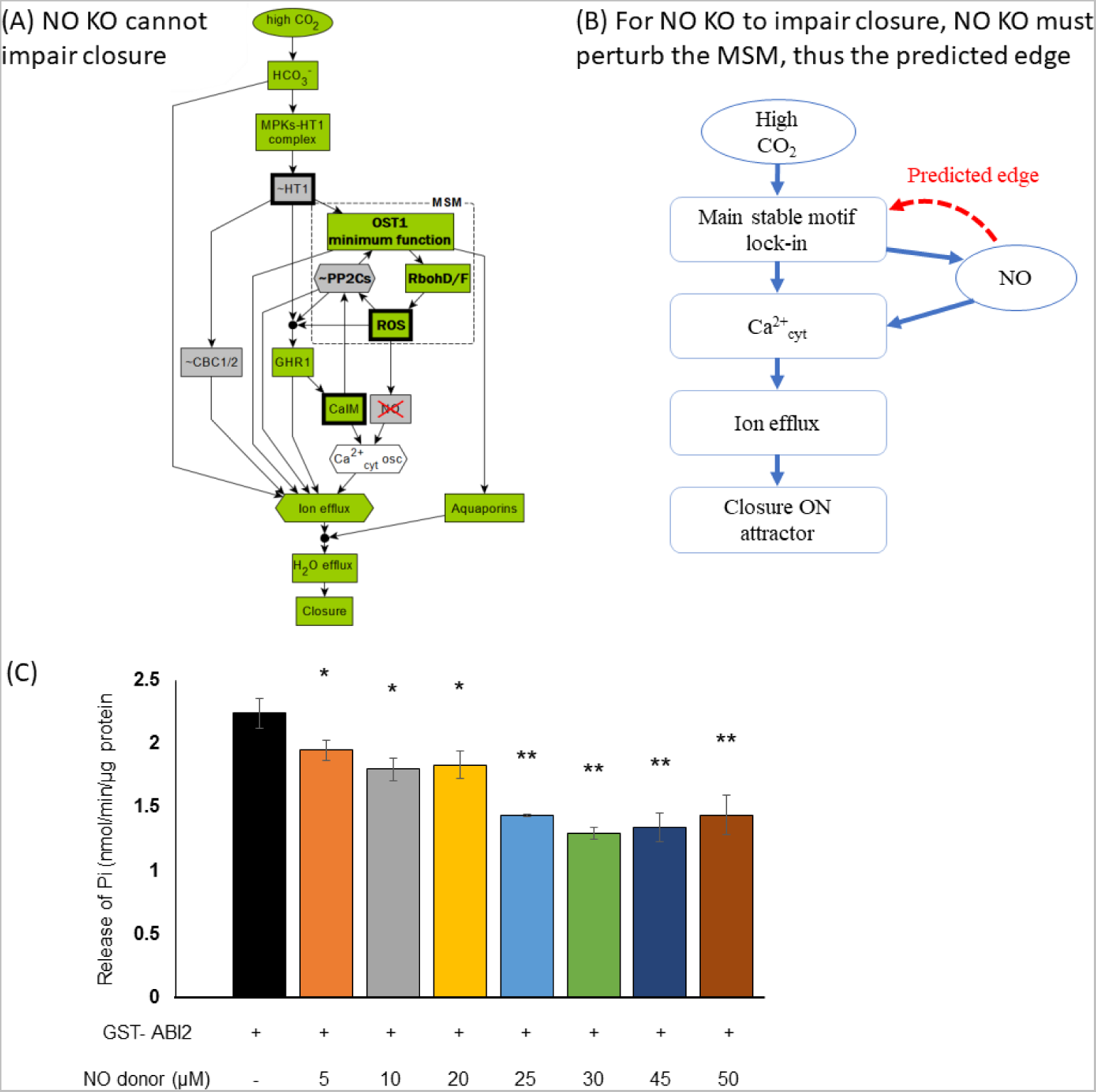
Network-based analysis predicted a new regulatory edge between NO and PP2Cs, which was validated by experiment. (A) Logic diagram showing that high CO_2_ can activate closure despite NO KO; (B) Conceptual sketch of NO’s role in the network and the predicted edge. NO must regulate the Main Stable Motif, otherwise loss of NO cannot impair high CO_2_ induced closure. (C) Experimental validation showing NO inhibits ABI2 phosphatase activity. The y axis shows the effect of increasing concentrations of the NO donor SNP on ABI2 activity, quantified by measuring the amount of phosphate released from a phosphopeptide substrate, as described in Methods. The bars indicate the means ± SE of three biological replicates. Significant differences between phosphatase activity of ABI2 alone and ABI2 treated with NO donor are indicated (**P < 0.001, *P < 0.05, Student’s t-test).

### Our model predicts which nodes of the high CO_2_ signaling network could be potential targets of currently unconnected nodes

There are signaling elements that are convincingly implicated in CO_2_ signaling, yet their connection to the signaling network remains unknown. During our literature review and data curation process, we placed such elements into the “insufficient connection” category. For example, *big1* mutants are compromised in elevated CO_2_-induced stomatal closure and bicarbonate activation of S-type anion channel currents [63], yet BIG1’s connection to known guard cell signaling elements remains undiscovered. Connecting these elements to the existing CO_2_ network can improve understanding of guard cell CO_2_ signaling and could lead to testable predictions. Our model offers a way to identify and prioritize potential targets of these unconnected elements based on the effects of node perturbations (Figure 6). The idea is that, if a node perturbation yields a similar response in the model to the observed effect of the unconnected signaling element, the node is then a potential target of the signaling element. For example, the *big1* mutant is known to have impaired CO_2_-induced closure. Our model simulations (Figure 6A) yield a comprehensive list of 24 single node perturbations that could impair closure. We predict that the *big1* mutant regulates one of these nodes; any alternative single target of *big1* would not cause loss of closure in the model. Moreover, one can utilize additional evidence to further narrow down the target candidates. For example, as *big1* mutants are also compromised in bicarbonate activation of S-type anion channel currents, we can further filter the 24 candidate nodes to those that satisfy both conditions: (1) the node is downstream of bicarbonate; and (2) the node’s perturbation results in reduced SLAC1 activity (reflected by the node AnionEM, which incorporates the effect of both SLAC1 channel nodes). This filter narrows the list of potential targets of BIG1 to 9 nodes, namely, MPKs, ABI1, ABI2, GHR1, HT1, OST1 minimum function, RbohD/F, and ROS. One can design experiments to test if some of these nodes are indeed perturbed in *big1* mutants. If a broader pool of candidate targets is needed, one could also consider the nodes whose perturbations causes hyposensitivity in high CO_2_ induced closure (five nodes in the last row of Table 3). These nodes may be appropriate because it might be difficult to distinguish reduced sensitivity and hyposensitivity in experiments.

**Table 3.** Model-predicted potential regulation targets to impair high CO_2_ induced closure. If a knockout of a signaling element impairs high CO_2_ induced closure, the model predicts the knockout will perturb one of these nodes. The first three rows are categorized based on the network location of the node, i.e., which component it is in. The last row is an extension where we hypothesize that perturbations causing hyposensitivity in the model could also be considered as having the potential to impair closure. Bold font indicates an example of 9 prioritized targets for BIG1, which are downstream of bicarbonate and whose perturbation can also cause reduced SLAC1 activity, as observed in the *big1* mutant.

| Category | Potential regulation target to impair closure |
| --- | --- |
| High CO<sub>2</sub> in-component or source node (6 nodes) | CA14, CO<sub>2</sub> porins, CO<sub>2</sub> entry of cell, HCO<sub>3</sub><sup>-</sup>, **MPKs, NADPH** |
| Signal transduction SCC (10 nodes) | **ABI1, ABI2, Ca<sup>2+</sup><sub>cyt</sub>, Ca<sup>2+</sup>ATPase, GHR1, HT1, OST1 minimum function, RbohD/F, ROS** |
| Ion Channel out-component (8 nodes) | H<sup>+</sup> ATPase, Anion EM, Aquaporins, Depolarization, H<sub>2</sub>O Efflux, K<sup>+</sup> efflux, KEV, KOUT |
| Nodes/perturbations causing hyposensitivity (5 nodes) | CaIM, CBC1/2, MPKs-HT1complex, QUAC1, SLAC1 CO<sub>2</sub> |

By a similar logic, the “hypersensitivity” response category in Figure 6A allows target prediction for signaling elements whose perturbation (knockout or constitutive activation) promotes closure. For example, faster closure is observed in plants lacking the ABC transporter AtABCB14 [64]. Hence, our model predicts that the knockout of AtABCB14 causes a node perturbation that leads to hypersensitivity, as annotated in Figure 6A. Furthermore, one can predict/prioritize targets of a signaling element as those that can induce closure in ambient CO_2_, based on the node perturbations that can include closure in ambient CO_2_ (Figure 6C). To summarize, our model provides ways to prioritize potential targets to link previously unconnected nodes with the high CO_2_ signaling network. Importantly, we only used BIG1 and AtABCB14 as examples; the proposed prediction method is general.

## III. Discussion

In this work we constructed a signaling network that integrates current information regarding the signaling mechanisms underlying high CO_2_ induced stomatal closure in guard cells. We assembled the nodes and edges of the network model from a thorough literature review process, with rigorous definitions of which signaling elements are involved in CO_2_ signaling and which are not; we also deployed a multi-node presentation to capture multiple mechanisms of the same element. Finally, we conducted a comprehensive model performance evaluation, making sure that our model indeed has high consistency with experimental observations. A previous model by Schroeder and colleagues of ABA and high CO_2_ induced closure, constructed as a case study of the Boolink graphical user interface [29], added to a model we originally created for ABA-induced stomatal closure [3] six nodes (CO_2_, CA1, CA4, MPK4/12, HT1, CBC1/CBC2) and changed five functions (GHR1, CaIM, Ca^2+^_c_, Microtubule, H_2_O efflux) without comprehensive evaluation of the model’s accuracy in recapitulating all experimental observations regarding high CO_2_ induced closure. In fact, two of the additions make the rest of the network superfluous: specifically, the function that controls CaIM has “OR CO_2_” in it, which means that high CO_2_ induces CaIM regardless of the rest of the regulators of CaIM; the function of H_2_O efflux also has “OR CO_2_” in it, which makes water efflux happen directly in response to CO_2_ regardless of the rest of the regulators of aquaporins. The combined effect of these two assumptions results in closure. In other words, the model of Karanam et al. short-circuits essentially the entire CO_2_ signaling network by making CO_2_ directly activate the very downstream part of the network. As a result, the model will not be able to capture the experimentally observed responses to many perturbations in the signaling network, e.g. the impaired high CO_2_ induced closure under *slac1*1 [47], *ht1-2* [10] and *mpk4/mpk12* [12] mutants. By comparison, a rigorous model construction process allows our CO_2_ model to comprehensively reproduce experimental observations related to high CO_2_ induced closure.

### Network methodologies reveal key decision-making motifs, pinpoint driver nodes, elucidate signal transduction mechanisms, predict new edges and responses

Our network-based modeling framework integrates piece-wise (node to node) biological knowledge to form a high-level comprehensive view of the signal transduction network underlying high CO_2_ induced stomatal closure. This framework complements conventional biological approaches that often focus on single regulators or single pathways. Our analysis supports recent findings that HT1 is a key decision-making node of the CO_2_ signaling system [13, 14], and offers additional insights regarding the ambient CO_2_ setting: closure induced by, e.g., exogenous ROS or Ca^2+^ in ambient CO_2_ must involve the inactivation of HT1 and the activation of a positive feedback loop (the Main Stable Motif, formed by ROS, OST1 and the PP2Cs). Conversely, under conditions that activate HT1 the nodes of this positive feedback loop may also lock-in to their opposite states and lead to closure OFF.

We have also introduced the concept of logic diagrams to visually depict the logical implications of events in the closure process, highlighting key nodes and pathways (subgraphs) responsible for decision-making of the system. The logic diagrams explain not only how signal transduction leads to stomatal closure under high CO_2_, but also how closure is lost or restored. Leveraging these concepts, we explain the mechanisms behind the five types of simulated closure responses in the presence of perturbations.

Combining these network-based methodologies allows the generation of model predictions that lead to novel experimental findings. It is important to note that the inconsistencies between of the model and experimental results also yield insights. For example, we found that the model is incapable of reproducing the experimental observation that NO knockout impairs high CO_2_ induced closure; by understanding that this inconsistency results from the NO-independent lock-in of the Main Stable Motif, we predicted that NO must regulate the MSM, and successfully validated this prediction experimentally, showing that NO inhibits ABI2, one of the PP2Cs in the MSM.

### Unexplained mechanisms in CO_2_ signaling

There are signaling elements known to mediate closure in response to a certain signal, but understudied in CO_2_ signaling; for example, PA (Phosphatidic acid). In our model construction process, when we designate each piece of knowledge regarding the regulation of a node by another into one of the five categories (see Supporting Info S1 and Table 1), we place such elements into the “generic”, or “insufficient evidence” categories. This way, our categorization of evidence can prioritize experiments to explore the roles of signaling elements in the context of CO_2_ signaling.

Our model construction and validation process included the examination of multiple different hypotheses regarding the role of certain nodes in high CO_2_ signaling. For example, the role of RHC1 in CO_2_ signaling is under debate: while Tian et al. showed experimental evidence that RHC1 is an essential CO_2_ signaling component [15], Tõldsepp et al. observed that a different *rhc1* mutant shows no reduction of high CO_2_-induced closure. We tested both versions of the HT1 regulatory function, with and without RHC1, and found that the function without RHC1 has a higher accuracy in capturing experimental observations. Therefore, we adopt the HT1 function without RHC1, more consistent with [16]. Our interpretation is that RHC1’s regulatory effect on HT1 is weaker than that of MPKs, thus it is insufficient to include in a Boolean model.

Some CO_2_ signaling mechanisms remain unclear. For example, observations about guard cell cytosolic pH suggests undiscovered buffering capacity in the cytosol. Brearley et al. and Xue et al. [8, 30] observed that cytosolic pH does not increase under high CO_2_ induced closure. On the other hand, after CO_2_ enters the guard cell, it undergoes the chemical reaction CO_2_ + H_2_O → HCO_3_^−^ + H^+^ catalyzed by CA1/4, producing H^+^. So, there must be some buffering capacity or other mechanism to keep cytosolic pH constant. Understanding this mechanism may affect other components in the network, and its refinement may improve the model as well as our understanding of the high CO_2_ signaling mechanism.

Future discovery of new signaling elements and interactions that participate in high CO_2_ induced closure will call for updates of this model, including the addition of new nodes and/or new edges, the possible removal of assumed edges (shown as dotted lines in Figure 1), or changes to the regulatory functions of certain nodes. It is important to note that even when new signaling elements and interactions are incorporated into the CO_2_ model, certain structural and dynamic features of the model will be preserved. For example, the network will still have a strongly connected component (see Figure 1B) that includes at least the 20 nodes currently in it. The updates will be such that the model preserves the current model’s agreements with experimental results. The coupling of the structural and dynamical preservation means that the decision-making role of the main stable motif will also be maintained in the updated model, even if newly added edges will increase its size (for example, by NO joining it). It is likely that newly discovered interactions will support the regulatory relationships that drive the decisions of the system. For example, the inhibition of ABI2 by NO is an additional mechanism by which NO contributes to the activation of the MSM. Ultimately, a cycle between experimental and modeling approaches offers the most effective way to elucidate the full network of high CO_2_ induced closure.

## IV. Methods and Data

### Construction of the CO_2_-signaling network and dynamic model

As described in the main text, we extract regulatory relationships from the literature (Supporting Info S1), and compile them into nodes and edges of the network (Supporting Info S2). Then we construct the regulatory functions (Supporting Info S3) based on the regulatory relationships.

### Simulation settings of the dynamical model

In simulation, we initialize the *in silico* stomata in a pre-stimulus open state, with closure-inhibiting nodes ON (namely ABI1, ABI2, HAB1, HT1, and H^+^ ATPase) and other nodes OFF (see SI S4 for complete initial condition). For example, Anion Efflux, a node that becomes activated during high CO_2_-induced closure, is initialized in the OFF state. We deploy a discrete-time simulation, with stochastic random order asynchronous update: at each discrete round of updates (or timestep), all nodes are updated in a randomly selected order. A different order is generated in each timestep. The main advantage of random order asynchronous update is that it can capture consensus dynamics without implementation of exact timing, as the actual timing and durations of the processes happening in the system are typically unknown. The stochasticity of the update helps avoid spurious behaviors that depend on synchrony (i.e., which only appear under synchronous update, where all nodes are updated at the same time, which is biologically unrealistic). We perform this simulation for 30 timesteps, long enough for the system to reach its attractor (stable long-time behavior). In this way, we can simulate how a node’s state changes over time during the stomatal closure process.

We perform 1000 replicate simulations, i.e., 1000 “*in silico* stomata” simulations, then calculate the “percentage of node activation” at each timestep as our metric, meaning the percentage of simulations in which the node (e.g. Closure) is in state 1/ON. For example, if 1000 replicate simulations yield 510 cases where closure is ON at time step 30, the simulated percentage of closure is 51%. In this way, oscillations and multistability with different node states will display a percentage of node activation value between 0% and 100%.

### Stable motif and attractor analysis

Stable motifs are the smallest strongly connected components in a system that can sustain a unique steady state for the constituent nodes [26, 55]. Stable motifs are identified in an expanded representation of the network (which includes the network’s regulatory functions as part of the network structure). A weaker version of a stable motif, called a conditionally stable motif [56], can sustain a unique steady state for its nodes as long as certain node(s) that regulate the motif are in a specific fixed state. The stable motifs and conditionally stable motifs of a Boolean network determine its attractors: one can uniquely associate sequences of stable motifs (stabilized in the order given by the sequence) to each attractor.

A driver (or driver set) of a stable motif is the node (or node combination) and its state that, when kept fixed, eventually leads to the locking in of the stable motif after sufficient updates. We used the python library pystablemotifs [27] to analyze the stable motifs, attractors and their control methods. The library is available through the github page https://github.com/jcrozum/pystablemotifs.

### Validation against experimental observations

We use the experimental observations in Supporting Info S1 that were not used in model construction (a subset of the indirect regulations, marked as “V” for validation in Supporting Info S1) as the ground truth for model validation. Because quantifying the (reduction of) stomatal aperture is difficult in both experiment and Boolean model, when comparing the experiment-based expected outcomes with the model simulation results, we employ a lenient criterion for consistency: any model-predicted defect (i.e. node percentage < 100%) is considered qualitatively consistent with either reduced response or no response. For example, if a simulation yields a closure percentage = 0.5, then it is considered consistent with both “reduced closure” and “no closure” in experimental observation. This type of consistency criterion was used in previous modeling as well [3]. We provide the comprehensive validation results in Supporting Info S4. The supplementary file also includes the trivially consistent cases due to circular reasoning, denote “true by construction”. For example, bicarbonate-induced activation of S-type anion channels was reduced in the dominant active *abi1* mutants (ABI1 CA) [43], but because ABI1 is a input in both SLAC1 functions and ABI1 ON is sufficient to deactivate both SLAC1 nodes, this experimental observation is captured by model/rule construction, and calling it a validation would be circular reasoning. This type of validation is not counted in our assessment of model accuracy performance.

### Definition of the response categories in CPC values, for simulated knockout or constitutive activity

We defined the cumulative percentage of closure (CPC) as the sum over 30 timesteps of the fraction of simulations that have the closure node in state 1 (ON), over 1000 replicate simulations. Thus, CPC ranges from 0 (if the percentage of closure were 0% at every time step) to 30 (if the percentage of closure were 100% at every time step). The CPC metric helps capture time-sensitive behavior such as hypo-/hyper-sensitivity, reflected as a small CPC difference from wildtype CPC.

We define the five categories of responses based on the CPC value and attractor state: close to wildtype, hypersensitivity, hyposensitivity, reduced closure, and no closure. We define the wildtype response range from the default high CO_2_-induced closure response. Since the simulation is stochastic (i.e. the CPC value can be different for each simulation), we determine the default CPC range by 600 batches of 1000 replicate simulations under the default high-CO_2_ signal, and define its CPC range as the “close to wildtype” CPC range (20.405 to 20.709). CPC values above this range are classified as hypersensitivity, meaning that the stomata reach the closure state significantly faster than wildtype. CPC values below the wildtype range with 100% percentage closure at the final timestep are classified as hyposensitivity. Reduced closure is the CPC value being much lower than wildtype range, with the final percentage of closure being between 0 and 1. In all of the observed cases in this last category, the closure value is oscillating in the attractor. Lastly, no closure is defined as the CPC value being zero or very close to zero, with final percentage of closure being zero. A small positive CPC can occur here, because closure may temporarily turn ON during the response process, though it cannot be maintained. The comprehensive CPC simulation results are provided in Supporting Info S6.

### Logic diagrams of the CO_2_ network model

The logic diagram is an integration of the network and node states that yields a specific outcome (e.g., the ON state of the node Closure). It is an expansion of the concepts of elementary signaling mode [57] and logical domain of influence [58], both of which are subgraphs of the expanded network (also called logical hypergraph), which incorporates the Boolean functions of all the nodes in the network [55]. The logic diagram indicates the node states visually: nodes whose name is preceded by ~ and have grey background are inactive, nodes with green background are active, and nodes whose name is followed by “osc” and have a white background oscillate. A single edge in a logic diagram represents a sufficient regulatory relationship that can achieve the indicated state of the target node; multiple edges incident on the same black dot represents “AND” logic, indicating that all nodes are required to achieve the target node state. A logic diagram may condense fully linear pathways of nodes and edges in the original network, to simplify visualization. This type of condensation does not change the regulatory logic of the network model [59].

To better help visual parsing in the CO_2_ network model, we also merged a few nodes of the original network. The node “PP2Cs” merges three members of the PP2C protein family, because these nodes have identical regulation. The technical definition of the merged node is PP2Cs = ABI1 or ABI2 or HAB1. The node “Ion flow” merges AnionEM and K^+^ efflux (i.e, Ion flow = AnionEM and K+ efflux). We also use a single node “Ca^2+^ osc” to represent the Ca^2+^ - Ca^2+^ ATPase negative feedback cycle, capturing the oscillating behavior of Ca^2+^ in the presence of a Ca^2+^ source (CaIM or CIS) and due to the inhibitory effect of the Ca^2+^ ATPase. We use hexagonal symbols for “PP2Cs” and “Ion efflux” as they represent a node set (Figure 5). Note the “Ion efflux” hexagon node needs multiple incident edges to activate, causing an exception of the edge definition.

The logic diagram can be read as a sequence of events and the conditions necessary for these events. For example, the sequence of events necessary for high CO_2_ induced closure described by Figure 5A is the following: high CO_2_ activates HCO_3_^−^, which inactivates HT1 via the formation of the MPKs-HT1 complex. HT1 being OFF then activate OST1 minimum function, which eventually activates the MSM. The simultaneous activation of the MSM and inactivation of HT1 (indicated by the black dot) activates GHR1, which in turn activates CPKs. GHR1 activates CaIM, which leads to cytosolic Ca^2+^ transients or oscillations. The MSM, GHR1, Ca^2+^_cyt_, HCO_3_^−^, as well as the inactivity of CBC1/2 contribute to the activation of Ion flows. The simultaneous activation of Ion flows and OST1-activated Aquaporin yield H_2_O efflux, which then leads to Closure. As another example, the logic diagram in Figure 6D shows that the activation of the MSM (e.g., by CaIM) leads to CIS, which then leads to cytosolic Ca^2+^ transients or oscillations, which activate MPK activity, which inhibits HT1, which then allows the activation of GHR1, then of the generic SLAC1 node, and finally the activation of the node Closure.

### Plant materials and growth conditions

*Arabidopsis thaliana* Columbia (Col-0) accession plants were used as wild-type. Col-0 were germinated on MS agar plates for 2 weeks and then grown in soil (1:1 mixture of Metro Mix 360: Sunshine Mix LC1 (Sun Gro Horticulture) in a growth chamber with an 8-h-light/16-h-dark cycle, 120 µmol photons m^−2^ s^−1^ light, and 20°C. Young fully expanded leaves from 4-5-week-old plants were used for stomatal aperture assay experiments.

### Stomatal aperture assay

Stomatal apertures were assayed as described previously with slight modifications [65]. In brief, epidermal peels were hand-peeled from the abaxial surfaces of leaves and then were floated interior-side down on an opening solution (20 mM KCl, 10 mM MES-KOH pH 6.15, 50 µM CaCl_2_) under white light (150 µmol m^−2^ sec^−1^) for 3 h to induce maximum opening of the stomata (initial aperture). After the 3h preincubation period, epidermal peels were floated in a six-well plate (each well with a diameter of 35 mm and height of 17 mm) containing pre-equilibrated high (800 ppm) or ambient (400 ppm) CO_2_ opening solution (approximately 3 mL). Subsequently, the multiwell cell culture plate was placed in a petri dish chamber and sealed with parafilm. The high or ambient CO_2_ concentration was continuously maintained within the chamber using a Licor-6400 (flow rate of 500 µmol s^-1^) while the samples were further incubated under white light (150 µmol m^−2^ sec^−1^). After 3 hr of incubation, the epidermal peels were imaged with a 40X objective using a light microscope (Nikon Diaphot 300) coupled to a digital camera (Nikon Coolpix 990). Stomatal apertures were measured in each image using ImageJ. To determine the effects of NO donor SNP (sodium nitroprusside dihydrate; Sigma 71778) on CO_2_-induced stomatal closure, experiments were performed according to the method described by [66] with some modifications. Briefly, epidermal peels were incubated for 3 hr in the opening buffer to induce stomatal opening. Subsequently, the epidermal peels were transferred to the opening buffer without or with SNP at the indicated concentrations and then treated under ambient CO_2_ (400 ppm CO_2_) or elevated CO_2_ (800 ppm CO_2_) in light for the indicated durations. A total of 30-40 stomatal apertures were measured per treatment for each individual experiment. Data presented are the mean of 3 independent experiments.

### Phosphopeptide-based phosphatase activity assay

To examine the effect of NO donor on ABI2 phosphatase activity, GST-tagged ABI2 was heterologously expressed in *Escherichia coli* and subsequently purified as described [67] with minor modifications. Briefly, pDEST15-ABI2 was constructed by transferring *ABI2* cDNA from pCR8/GW/TOPO TA vector to the N-terminal GST fusion expression vector pDEST15 (Invitrogen) by Gateway LR recombination (Invitrogen). The cloned construct was verified by sequencing. The GST-ABI2 protein was expressed in *E. coli* BL21(DE3) pLysS Rosetta cells by induction with 0.5 mM IPTG for 16 hr at 16 °C. The cells were harvested by centrifugation (6000 g, 8 min, 4°C) and stored at −80°C until use. All purification procedures were carried out at 4 °C. The final cell pellet was resuspended in 10 ml of B-PER reagent (Pierce, catalog number 78243) supplemented with 1 mg/ml lysozyme (Sigma, catalog number L6876), 25 mg/ml DNAse and EDTA-free complete protease inhibitor (Thermo Fisher scientific, catalog number A32955) and incubated for 60 min at 4 °C with gentle shaking. The cell lysate was spun at 16000g for 15 min. at 4 °C and the resulting supernatant was filtered using a 0.45 µm filter. Subsequently, the supernatant was loaded onto a 5 ml QIAGEN polypropylene column packed with GST glutathione agarose resin (Pierce, 16100) and the column-bound protein was eluted with equilibration buffer (25 mM Tris-HCl pH 7.5, 150 mM NaCl) supplemented with 20 mM reduced L-glutathione (Sigma, catalog number G4251). The purity of the recombinant GST-ABI2 was estimated by analysis of aliquots separated on a 12% SDS polyacrylamide gel stained with Gel-Code Blue (Thermo), and the concentration was estimated based on known concentrations of Fraction V BSA (Pierce, catalog number 23209) run on the same gel.

The effect of NO donor on ABI2 phosphatase activity was carried out using the Ser/Thr phosphatase assay kit (Promega; catalog number V2460). The reaction was performed in a 50 µl volume containing 28 nM of purified recombinant GST-ABI2 protein in the presence or absence of the indicated concentration of NO donor SNP. After 30 min incubation at 30 °C, the reaction was stopped by addition of 50 µl of molybdate dye/additive solution. The resulting mix was incubated for another 20 min at room temperature followed by absorbance measurements at 630 nm in a Synergy Neo2 multimode plate reader (Biotek). The amount of released phosphate was calculated based on a standard curve obtained using phosphate standard solutions of known concentration.

## Acknowledgements

The authors thank Dr. David Chakravorty, Dr. Yotam Zait, Dr. Ángel Ferrero-Serrano, and Dr. Jordan C. Rozum for helpful discussions.

## Supporting Information files

S1 Table. Summary of evidence from our literature review for CO_2_ network and model construction.

S2 Table. Description of the 47 nodes and 95 edges of the CO_2_ network

S3 Text. Regulatory functions of the dynamic model and their justifications

S4 Text. Supporting Information on signaling elements whose participation in CO_2_ signaling has not been tested and on minor conditionally stable motifs in the CO_2_ network.

S5 Table. Comparison of the model simulation results with experimental observations

S6 Table. CPC results of systematic perturbation (KO/CA) simulations under high CO_2_ and ambient CO_2_

## Competing interests

The authors declare that no competing interests exist. The authors do not have a related or duplicate manuscript under consideration (or accepted) for publication elsewhere. This study does not involve human or animal research ethics.

## Financial Disclosure Statement

This work was supported by NSF grant MCB 1715826 to S.M.A. and R.A. Link: https://www.nsf.gov/awardsearch/showAward?AWD_ID=1715826. The funders had no role in study design, data collection and analysis, decision to publish, or preparation of the manuscript.

## Data Availability

Data in this study are provided in the Supporting Information files.

## Author contributions

Conceptualization: Xiao Gan, Reka Albert, Sarah M. Assmann.

Data curation: Xiao Gan, with input from all authors.

Formal analysis: Xiao Gan, Kyu Hyong Park.

Funding acquisition: Reka Albert, Sarah M. Assmann.

Investigation: All authors.

Methodology: Xiao Gan, Palanivelu Sengottaiyan, Kyu Hyong Park, Reka Albert.

Project administration: Reka Albert, Sarah M. Assmann.

Software: Xiao Gan, Kyu Hyong Park.

Supervision: Reka Albert, Sarah M. Assmann.

Validation: Palanivelu Sengottaiyan, Xiao Gan.

Visualization: All authors.

Writing: Xiao Gan, Reka Albert, and Sarah M. Assmann, with input from all authors.

---

**Table 1.** Edge list of the high CO_2_ induced closure network. The edges represent interactions and regulatory effects. The start node (first column) indicates the source of the interaction and the end node (second column) indicates the target of the interaction. “Type” records the sign and type of the edge. Positive edge sign, “+”, indicates positive/activating regulation and “–” indicates negative/inhibitory regulation. For edge type, we use “D” for direct regulation, “I” for indirect regulation, and “A” for assumed regulation. In rare cases we use “D/A” to represent an edge that is supported by a direct interaction but has assumed properties, e.g. assumed direction or sign. “Ref.” column records the reference that supports the edge (“P” is short for “Palani’s preprint”). Edges representing known biophysical/biochemical processes, e.g. AnionEM → Depolarization, do not have a reference. The table follows alphabetic order of the start node.

| Start node | End node | Type | Ref. |
| --- | --- | --- | --- |
| 8-nitro-cGMP | ADPRc | +, A | [68-70] |
| ABI1 | OST1 minimum function | -, D | [71-73] |
| ABI1 | OST1 activated | -, D | [71-73] |
| ABI1 | SLAC1 generic | -, I | [74-76] |
| ABI1 | SLAC1 CO<sub>2</sub> | -, I | [74-76] |
| ABI2 | OST1 minimum function | -, D | [73] |
| ABI2 | OST1 activated | -, D | [73] |
| ABI2 | GHR1 | -, D | [50] |
| ABI2 | SLAC1 generic | -, I | [74-76] |
| ABI2 | SLAC1 CO<sub>2</sub> | -, I | [74-76] |
| ADPRc | cADPR | +, D |  |
| AnionEM | Depolarization | +, D |  |
| AnionEM | H<sub>2</sub>O Efflux | +, D |  |
| Aquaporins | H<sub>2</sub>O Efflux | +, D | [6, 77] |
| Ca<sup>2+</sup> ATPase | Ca<sup>2+</sup><sub>cyt</sub> | -, D |  |
| CA1/4 | HCO<sub>3</sub><sup>-</sup> | +, D | [7, 8] |
| CA1/4 | RHC1 | +, D | [15] |
| Ca<sup>2+</sup><sub>cyt</sub> | MPK activity | +, I | [11, 35, 78] |
| Ca<sup>2+</sup><sub>cyt</sub> | Ca<sup>2+</sup> ATPase | +, D | [23] |
| Ca<sup>2+</sup><sub>cyt</sub> | CPKs | +, D | [75] |
| Ca<sup>2+</sup><sub>cyt</sub> | QUAC1 | +, A | [79] |
| Ca<sup>2+</sup><sub>cyt</sub> | H<sup>+</sup> ATPase | -, D | [80] |
| Ca<sup>2+</sup><sub>cyt</sub> | KEV | +, D | [81] |
| Ca<sup>2+</sup><sub>cyt</sub> | Depolarization | +, D |  |
| cADPR | CIS | +, I | [82] |
| CaIM | ABI1 | -, I | [61] |
| CaIM | ABI2 | -, I | [61] |
| CaIM | HAB1 | -, I | [61] |
| CaIM | Ca<sup>2+</sup><sub>cyt</sub> | +, D |  |
| CBC1/2 | SLAC1 CO<sub>2</sub> | - | [83] |
| cGMP | 8-nitro-cGMP | +, D |  |
| CIS | Ca<sup>2+</sup><sub>cyt</sub> | +, D |  |
| CO<sub>2</sub> entry of cell | HCO<sub>3</sub><sup>-</sup> | +, D | [7, 8] |
| CO<sub>2</sub> porins | CO<sub>2</sub> entry of cell | +, D | [6] |
| CPKs | CPKs | +, D/A | [84] |
| CPKs | SLAC1 generic | +, D | [75, 76, 85] |
| Depolarization | KOUT | +, D |  |
| GHR1 | CaIM | +, A | [50] |
| GHR1 | CPKs | +, D | [41] |
| GHR1 | SLAC1 generic | -, D | [50] |
| GHR1 | SLAC1 CO<sub>2</sub> | -, D | [50] |
| GTP | cGMP | +, D | [86] |
| H<sup>+</sup> ATPase | Depolarization | -, D |  |
| H<sub>2</sub>O Efflux | Closure | +, D |  |
| HAB1 | OST1 minimum function | -, D | [73] |
| HAB1 | OST1 activated | -, D | [73] |
| HCO<sub>3</sub><sup>-</sup> | MPKs-HT1 complex | +, D | [11-14, 16] |
| HCO<sub>3</sub><sup>-</sup> | RHC1 | +, D | [15] |
| HCO<sub>3</sub><sup>-</sup> | SLAC1 CO<sub>2</sub> | +, A | [36] |
| High CO<sub>2</sub> | CO<sub>2</sub> entry of cell | +, D |  |
| HT1 | CBC1/2 | +, D | [13] |
| HT1 | GHR1 | -, D | [11, 15] |
| HT1 | OST1 minimum function | -, D | [11, 15] |
| HT1 | OST1 activated | -, D | [11, 15] |
| K<sup>+</sup> efflux | Depolarization | -, D |  |
| K<sup>+</sup> efflux | H<sub>2</sub>O Efflux | +, D |  |
| K<sup>+</sup> efflux | K<sup>+</sup> efflux | +, A |  |
| KEV | Depolarization | +, D |  |
| KEV | K<sup>+</sup> efflux | +, D |  |
| KOUT | K<sup>+</sup> efflux | +, D |  |
| MPKs | MPKs-HT1 complex | +, D |  |
| MPKs | MPK activity | +, D |  |
| MPKs-HT1 complex | HT1 | -, D | [11-14] |
| MPK activity | HT1 | -, A | [11, 16, 35] |
| MPK activity | MPK activity | +, D/A | [87] |
| MPK activity | SLAC1 generic | +, A | [35] |
| NAD<sup>+</sup> | cADPR | +, D |  |
| NADPH | NO | +, D | [51] |
| NADPH | ROS | +, D |  |
| NIA1/2 | NO | +, D | [51] |
| Nitrite | NO | +, D | [51] |
| NO | 8-nitro-cGMP | +, I | [53] |
| NO | NOGC | +, D | [86] |
| NO | KOUT | -, I | [88] |
| NOGC | cGMP | +, D | [86] |
| OST1 activated | SLAC1 generic | +, D | [74] |
| OST1 minimum function | OST1 activated | +, D |  |
| OST1 minimum function | RbohD/F | +, D | [89] |
| OST1 minimum function | SLAC1 generic | +, D | [74, 90] |
| OST1 minimum function | SLAC1 CO<sub>2</sub> | +, D | [74, 90] |
| OST1 minimum function | QUAC1 | +, D | [91] |
| OST1 minimum function | Aquaporins | +, D | [77] |
| QUAC1 | AnionEM | +, D | [45] |
| RbohD/F | ROS | +, D |  |
| RHC1 | HT1 | -, D | [15] |
| ROS | GHR1 | +, I | [50] |
| ROS | ABI1 | -, D | [92] |
| ROS | ABI2 | -, D | [93] |
| ROS | HAB1 | -, D | [94] |
| ROS | 8-nitro-cGMP | +, I | [53] |
| ROS | NIA1/2 | +, A | [95] |
| ROS | H<sup>+</sup> ATPase | -, D | [96] |
| ROS | KOUT | -, I | [97] |
| SLAC1 CO<sub>2</sub> | AnionEM | +, D |  |
| SLAC1 nonCO<sub>2</sub> | AnionEM | +, D |  |

## Notes

### Competing Interest Statement

The authors have declared no competing interest.

